# Body size as a magic trait in two plant-feeding insect species

**DOI:** 10.1101/2022.10.11.511791

**Authors:** Ashleigh N. Glover, Emily E. Bendall, John W. Terbot, Nicole Payne, Avery Webb, Ashley Filbeck, Gavin Norman, Catherine R. Linnen

## Abstract

When gene flow accompanies speciation, recombination can decouple divergently selected loci and loci conferring reproductive isolation. This barrier to sympatric divergence disappears when assortative mating and disruptive selection involve the same “magic” trait. Although magic traits could be widespread, the relative importance of different types of magic traits to speciation remains unclear. Because body size frequently contributes to host adaptation and assortative mating in plant-feeding insects, we evaluated several magic trait predictions for this trait in a pair of sympatric *Neodiprion* sawfly species adapted to different pine hosts. A large morphological dataset revealed that sawfly adults from populations and species that use thicker-needled pines are consistently larger than those that use thinner needled-pines. Fitness data from recombinant backcross females revealed that egg size is under divergent selection between the preferred pines. Lastly, mating assays revealed strong size-assortative mating within and between species in three different crosses, with the strongest prezygotic isolation between populations that have the greatest interspecific size differences. Together, our data support body size as a magic trait in pine sawflies and possibly many other plant-feeding insects. Our work also demonstrates how intraspecific variation in morphology and ecology can cause geographic variation in the strength of prezygotic isolation.

## Introduction

A core goal of evolutionary biology is to understand how new species evolve via the accumulation of reproductive barriers (Coyne and Orr 2004; Nosil 2012; Butlin et al. 2012; Abbott et al. 2013; Hernández-Hernández et al. 2021). Historically, the evolution of reproductive isolation was thought to require geographic barriers that prevent gene flow and recombination from breaking apart associations between loci under divergent selection and loci responsible for non-random mating, both of which are required for coexistence in sympatry (Mayr 1963; Futuyma and Mayer 1980; Felsenstein 1981). Over the last two decades, however, theoretical and empirical studies have demonstrated that sympatric speciation (the evolution of reproductive isolation in the absence of barriers to gene exchange) can and does occur, although the prevalence is still debated (Berlocher and Feder 2002; Gavrilets 2003; Bolnick and Fitzpatrick 2007; Foote 2018). One mechanism for overcoming the “selection-recombination antagonism” in divergence-with-gene-flow scenarios is pleiotropy: if a trait under divergent selection and a trait involved in non-random mating are encoded by the same gene, recombination cannot decouple the two (Gavrilets 2004; Servedio et al. 2011; Thibert-Plante and Gavrilets 2013). Such traits, which have been dubbed “magic traits” (Gavrilets 2004), can facilitate sympatric speciation (Dieckmann and Doebeli 1999) and are potentially widespread in nature (Servedio et al. 2011). However, the importance of magic traits versus other mechanisms that promote speciation-with-gene-flow (e.g., physical linkage, Kirkpatrick and Barton 2006; Ravinet et al. 2017) and the relative importance of different types of magic traits are currently unknown (Servedio et al. 2011).

Plant-feeding insects are classic models of magic traits and sympatric speciation via host shifts (Bush 1975; Berlocher and Feder 2002; Drès and Mallet 2002; Matsubayashi et al. 2010). Because herbivorous insects often rely on their host plant for much (or all) of their life cycle, colonization of a new host plant can result in strong selection on traits required to locate, feed, and reproduce on the new host (Forbes et al. 2017). When different traits are favored on different hosts, reproductive isolation can evolve between host-associated populations as a by-product of divergent selection (via pleiotropy or linkage) or via direct selection for reduced gene exchange (e.g., reinforcement, Berlocher and Feder 2002; Matsubayashi et al. 2010; Nosil 2012). Because many plant-feeding insects mate on their host plant and divergent selection between hosts can involve many traits (Via and Hawthorne 2002; Nosil and Sandoval 2008; Matsubayashi et al. 2010), this life history may be especially conducive to producing magic traits. Indeed, there are numerous examples of magic traits in plant-feeding insects, including habitat choice in *Rhagoletis pomonella* apple maggot flies (Feder et al. 1994; Feder 1998), phenology in *Ostrinia nubilalis* European corn borer moths (Dopman et al. 2005; Levy et al. 2015; Kozak et al. 2019; Kunerth et al. 2022), wing color patterning in *Heliconius* butterflies (Jiggins et al. 2006; Merrill et al. 2012; Mullen and Shaw 2014), body color in *Timema cristiniae* stick insects (Nosil et al. 2002; Greenfield 2016), and signal frequency in *Enchenopa binotata* treehoppers (Cocroft et al. 2008). By comparison, and despite numerous vertebrate examples (Servedio et al. 2011), few studies have evaluated body size as a magic trait in plant-feeding insects (but see Augustyn et al. 2017).

As is the case in many non-insect taxa (Blackburn et al. 1999; Ólafsdóttir et al. 2006; Liu et al. 2018; Stobo-Wilson et al. 2020), body size is highly variable both within and between insect species (Chown and Gaston 2010; Beukeboom 2018; Tseng and Pari 2019) and subject to both sexual selection and natural selection via multiple biotic and abiotic agents (Common et al. 2020; Dudaniec et al. 2022). In herbivorous insects, body size is often constrained by physical or visual properties of the host plant (Matsubayashi et al. 2010). For example, immature stages may need to fit within host tissue as eggs or feeding larvae and adult females may need appropriately sized ovipositors for successful egg laying (Bendall et al. 2017). Having the correct body size can also be important for securely attaching to the host plant or effective crypsis to evade predators (Augustyn et al. 2017). In addition, compatible female and male body sizes are important for proper alignment during mating in many insect species (e.g., Weissman et al. 2008; Villa et al. 2019). This may explain, in part, why size-assortative mating is widespread in insects (Bonduriansky 2001). Given its potentially widespread role in both divergent host adaptation and assortative mating, body size could be an unusually common magic trait in plant-feeding insects. As a first step to evaluating this possibility, we investigated body size, host adaptation, and assortative mating in a sister-species pair of pine sawflies, *Neodiprion lecontei* and *Neodiprion pinetum*.

*Neodiprion* (Order: Hymenoptera; Family: Diprionidae) are pine-feeding sawflies that depend on their host plant, mostly pines in the genus *Pinus*, at all stages of the life cycle: the adults mate on the host; adult females lay their eggs in the needle tissue; the larvae feed on the needles; cocoons are spun directly on or underneath the host tree (Coppel and Benjamin 1965; Knerer and Atwood 1973). *Neodiprion* sawflies have long been recognized as having traits conducive to sympatric speciation (Knerer and Atwood 1973; Bush 1975; Strong et al. 1984; Linnen and Farrell 2010), and genetic data indicate that many species diverged with gene flow (Linnen and Farrell 2007). Within this genus, sister species *N. lecontei* and *N. pinetum* have large, mostly overlapping ranges (Fig. 1A; Linnen and Farrell 2008, 2010) and, for several reasons, are a good model for studying speciation-with-gene-flow and magic traits. First, a recent demographic analysis supports a sympatric divergence scenario for *N. lecontei* and *N. pinetum* (Bendall et al. 2022). Second, *N. lecontei* and *N. pinetum* are not completely reproductively isolated: viable and fertile hybrids can be produced in the lab (Bendall et al. 2017, 2021) and these species are known to hybridize in some parts of their ranges (Bendall et al. 2022). Incomplete isolation is essential for distinguishing between reproductive barriers that cause speciation versus those that accrue only after it is complete (Coyne and Orr 2004; Nosil 2012). Lastly, these species are adapted to pine hosts with very different needle sizes. *N. pinetum* only uses white pine (*P. strobus*) and is the only species in the genus that prefers this thin-needled host. *N. lecontei* avoids white pine and uses other *Pinus* species that have thicker needles (Wilson et al. 1992; Linnen and Farrell 2010; Bendall et al. 2017). Because female *Neodiprion* must embed their eggs within the needles of their host pine, *N. pinetum* and *N. lecontei* have evolved divergent egg-laying traits. Compared to *N. lecontei*, the white pine specialist (*N. pinetum*) has a much stronger preference for laying eggs in the thin-needled host, as well as a thinner, shorter, and straighter ovipositor (the saw-like appendage used to cut out egg pockets in the needle tissue; Fig. 1B) and a tendency to lay fewer eggs per needle (Bendall et al. 2017). Overall, previous data point to a sympatric divergence scenario with the potential for strong divergent selection on body size stemming from differences in host needle size.

**Figure 1.**
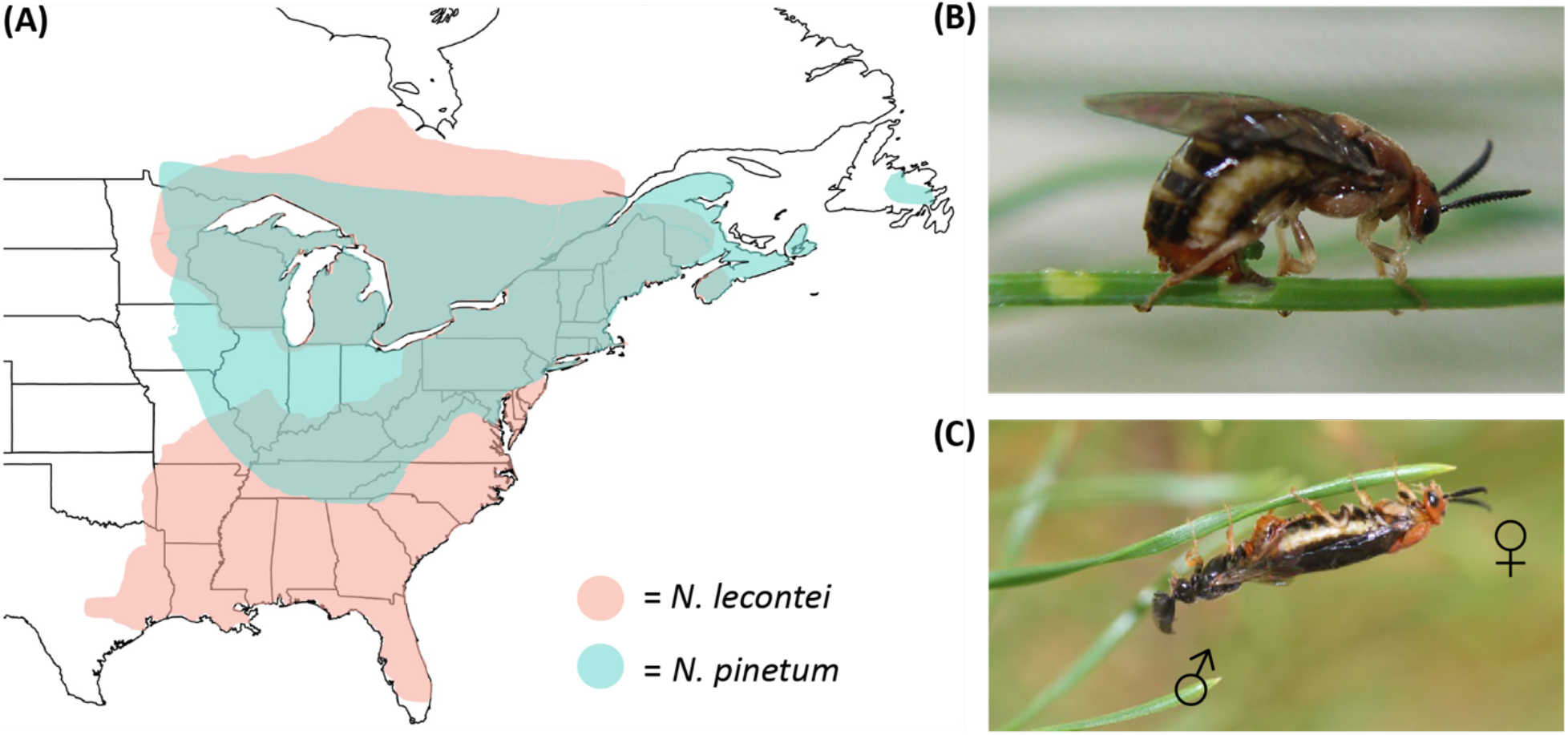
Rationale for body size as a magic trait in two largely sympatric pine sawfly species. (A) Approximate geographic ranges of *Neodiprion lecontei* (coral shading) and *N. pinetum* (teal shading). Note that most of *N. pinetum*’s range is contained within *N. lecontei*’s range. (B) A *N. pinetum* female uses her saw-like ovipositor to cut egg pockets into her thin-needled white pine host (photo by R.K. Bagley). Because eggs and ovipositors must fit within pine needles, differences in host needle width may act as a source of divergent selection on *Neodiprion* body size. (C) Mating pair of *N. lecontei* adults, with adult male on the left (photo by R.K. Bagley); for video, see Video S1. Because matched female and male body sizes are likely important for maintaining proper alignment during mating, body size could be an important source of assortative mating in *Neodiprion*.

Here, we evaluate several magic trait predictions for body size in *N. lecontei* and *N. pinetum*. First, for body size to be a magic trait, it must differ between *N. lecontei* and *N. pinetum*. Because needle width varies both within and between *Pinus* species (Nobis et al. 2012; Bendall et al. 2017), we also explore the potential for spatial variation in body size selection related to needle width. We predict both that geographic variation in adult body size for each species will correspond to geographic variation in host needle width and that *N. lecontei* adults will be larger than *N. pinetum* adults overall. Second, to be a magic trait, body size must be under divergent selection. For successful development, *Neodiprion* eggs must be completely embedded in pine needles, which requires that egg size is well matched to the width of the pine needles. Because *N. lecontei* and *N. pinetum* females prefer to lay eggs in thick-needled and thin-needled pines respectively (Bendall et al. 2017), we hypothesize that there is a trade-off between egg provisioning (bigger eggs produce bigger larvae with higher survival; Fox and Czesak 2000; Macke et al. 2011) and egg fitting (if eggs are too big for a needle, they will desiccate and die). This trade-off should favor larger eggs in *N. lecontei* and smaller eggs in *N. pinetum*. As larger eggs tend to produce larger larvae and adults in Hymenoptera (Fox and Czesak 2000; Macke et al. 2011; Church et al. 2019) and other insects (Fox and Czesak 2000; Fischer et al. 2002; Meister et al. 2018), divergent selection on egg size would produce differently sized adults of both sexes. Thus, we predict that *N. pinetum* will have smaller eggs and that smaller eggs will increase hatching success on the thin-needled white pine, while larger eggs will increase hatching success on thicker-needled pines. Lastly, to be a magic trait, body size not only has to be under divergent selection, but also must be involved in assortative mating. In pine sawflies, successful mating requires proper body alignment as males attempt to establish a tight genital connection with females that often exhibit resistance behaviors (Fig. 1C and Video S1). Thus, we predict that both species will exhibit size-assortative mating and that this will contribute to prezygotic isolation between them. To evaluate these predictions, we combine morphological data from pine sawflies and their hosts with fitness data from recombinant backcross females and no-choice mating assays. Our data are consistent with the hypothesis that body size is a magic trait in *Neodiprion* sawflies, raising the possibility that body size divergence may play a more important role in plant-feeding insect speciation than has been previously appreciated.

## Materials and Methods

### Geographic variation in host needle width and adult body size

Because *Neodiprion* eggs must be completely embedded in pockets that adult females carve within pine needles, egg size and female ovipositor size are likely constrained by needle width (Bendall et al. 2017). Given these constraints, we predicted that geographic variation in adult body size would correlate with geographic variation in host needle width. Although the ideal way to test this prediction would have been to collect host needles and sawfly adults from the same sites, we only had access to pine and sawfly samples that were collected at different times and without this hypothesis in mind. We therefore analyzed geographic variation in host needle width and adult body size separately, with the following expectations: *N. lecontei* adults and host needles will be bigger than those of *N. pinetum* and geographic trends in body size would mirror those observed for needle width (i.e., have slopes with the same sign and similar magnitude).

To characterize geographic variation in pine needle width, we sampled needles from multiple trees and sites for each of 10 *Pinus* species used by *N. lecontei* (*P. taeda, P. palustris, P. echinata, P. elliottii, P. clausa, P. glabra, P. virginiana, P. rigida, P. resinosa*, and *P. banksiana*) and the single host species used by *N. pinetum* (*P. strobus*). For each pine species, we collected clippings from 60-100 trees sampled from 6-10 sites (9-10 trees per site). Using digital calipers (Mitutoyo CD-6’PMX), we measured the width of 3 randomly sampled needles per tree and averaged these measurements to obtain a single needle width value per tree. In total, we sampled 878 individual pine trees from 88 sites (Table S1). To determine how needle width varies as a function of host species and latitude, we used linear regression to model needle width as a function of pine species, latitude, and their interaction. We used a Type III Analysis of Variance (ANOVA), implemented in the *car* v3.1-0 package, to evaluate the significance of model terms and the *emmeans* v1.8.0 package for post-hoc tests with false discovery rate (FDR) correction for multiple testing. These and all other statistical analyses were performed in R version 4.1.0 (R Core Team, 2021). Because we were specifically interested in comparing body size clines between sawflies with needle width clines from their pine hosts, we also used linear regression to estimate the relationship between (1) needle width and latitude for all *N. lecontei* hosts (regardless of host species) and (2) needle width and latitude for *P. strobus*, the only *N. pinetum* host pine.

To characterize geographic variation in body size for adult females and males of both species, we collected *N. lecontei* and *N. pinetum* across the eastern United States between 2015 and 2021 (Table S2 and Fig. S1) and reared immature stages to adults in the lab using host plant clippings and standard lab protocols (Harper et al. 2016; Bendall et al. 2017). Upon emergence, live adults (which are non-feeding) were either preserved immediately or stored at 4°C to prolong life until needed for propagating lab lines or use in experimental assays. All adults were ultimately placed in 100% ethanol and stored at -20°C. We used a Neiko Tools Digital Caliper (model 01407A) to measure body length (i.e., from the tip of the head to the tip of the abdomen) in millimeters (mm) for 1,080 preserved adults (*N. lecontei* females: *N* = 328; *N. lecontei* males: *N* = 243; *N. pinetum* females: *N* = 279; *N. pinetum* males: *N* = 230; Table S2). To determine how body length differed as a function of latitude, species, and sex, we fit a linear model to the body size data, with latitude, species, sex, latitude x species, latitude x sex, and species x sex as fixed effects. We used a Type III ANOVA to assess significance of model terms. Based on these results, we also fit individual geographic clines to each sex for each species.

### Divergent selection on egg size

We hypothesize that variation in needle width among pine populations and species generates divergent selection on body size (egg size) in *Neodiprion* that use different pine hosts via a combination of constraints imposed by thin needles (favors smaller eggs) and selection on early larval survival (favors larger eggs). To test this hypothesis, we first verified that *N. pinetum* (thin-needled specialist) and *N. lecontei* (uses hosts with thicker needles) differ in egg size. To do so, we used females reared from wild-caught larvae collected from different sites in Kentucky and Tennessee between 2013 and 2015 (Table S3). In *Neodiprion*, all egg maturation occurs within the cocoon, from which females emerge with a full complement of eggs ready for oviposition (i.e., these species are pro-ovigenic). To measure egg length and width, we dissected eggs out of the abdomens of recently eclosed adult females and photographed a random subset of eggs at 10X total magnification using a Zeiss Discovery V8 stereomicroscope with an Axiocam 105 color camera and ZEN lite 2012 software (Carl Zeiss Microscopy, LLC Thornwood, NY). We then used the ZEN lite software to measure the length and width of 5 eggs from each of 5 females from each species (*N* = 25 eggs per species). We then calculated egg area after García-Barros (2000) using the following equation:

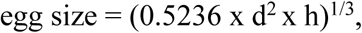

where d = egg diameter (width) and h = egg length. To test for differences in egg size between *N. pinetum* and *N. lecontei*, we fit a linear mixed-effects model to the egg area data, with species as a fixed effect and female ID as a random effect (since multiple eggs were sampled from each female). To evaluate significance of the species term, we used a Type II ANOVA.

*N. lecontei* and *N. pinetum* differ in many traits that affect hatching success on different types of pines, including ovipositor size and number of eggs per needle (Bendall et al. 2017; Fig. 3A, 3B). To evaluate the effect of egg size independent of other traits, we used a backcross design to generate recombinant hybrid progeny between *N. lecontei* and *N. pinetum* after Bendall et al. (2017). All females were *lecontei*-backcross females (one set of *N. lecontei* chromosomes and one set of chromosomes with approximately 50% *N. lecontei* and 50% *N. pinetum* ancestry). We used a *lecontei*-backcross design because preference for the thin-needled *N. pinetum* host is dominant (Bendall et al. 2017), and we wanted to ensure a good sample size of females that laid on both hosts. Backcross females were released individually into mesh cages (60 × 40 × 40 cm) containing two seedlings of *Pinus banksiana* (*N. lecontei* host) and two seedlings of *P. strobus* (*N. pinetum* host). Cages were checked daily for oviposition and once this occurred, live females (which tend to remain at the bottom of the egg-bearing branch until death) were preserved in 100% ethanol for subsequent morphological and molecular work. After preserving females, the number of eggs laid on each seedling were counted and monitored daily for hatching. Because females almost always cluster their eggs in a single branch tip, host choice was scored as a binary trait (*P. strobus* or *P. banksiana*). Once a hatchling was observed, eggs were given an additional 48 hours to provide sufficient time for hatching. All hatchlings were then removed via washing them off the pine seedling in ethanol. Hatchlings were counted and the hatching success for each female was calculated as the proportion of eggs that hatched (number of hatchlings/number of eggs).

After oviposition, females almost always have eggs left in the abdomen. We used these remaining eggs to quantify average egg size for each female. To measure egg size, we first rehydrated preserved female abdomens by soaking them in five decreasing concentrations of ethanol (95%, 80%, 65%, 50%, and 25%) for 10 minutes each. After the lowest ethanol concentration, we soaked females in deionized water for 24 hours at room temperature. We then dissected the eggs from each female’s abdomen and placed them in 300 μL of a modified Ringer’s dissection solution (7.5 g/L NaCl, 0.35 g/L KCl, and 0.21 g/L MgCl_2_). We then imaged 1-10 eggs per female at 10X total magnification and measured eggs and calculated area as described above. We then averaged these values to obtain a single average egg area per female. Although ethanol preservation and rehydration may cause preserved eggs to differ in size from fresh eggs, this approach should nevertheless reveal *relative* egg size differences among females (i.e., which females laid the largest or smallest eggs) because all eggs were treated in the same way. We are also assuming here that the size of leftover eggs correlates positively with the size of laid eggs.

In total, we measured hatching success, egg size, and host preference for *N* = 38 *lecontei*-backcross females. After removing two females that laid eggs on both pine species, our final sample size was *N* = 36 backcross females. To evaluate the effect of egg size (egg area) and host (*P. banksiana* or *P. strobus*) on hatching success, we used the *glm* function (*lmerTest* v3.1-3) to fit a mixed-effects logistic regression model to the hatching data (proportion hatched ∼ egg area + host + area*host), followed by a Type III ANOVA to evaluate the significance of model terms. A significant egg area x host interaction would support our hypothesis that there is a trade-off between egg fit and egg provisioning.

### Size-assortative mating and reproductive isolation

To determine whether *Neodiprion* adults mate assortatively by size and to quantify reproductive isolation between *N. pinetum* and *N. lecontei*, we first sampled larval colonies of both species from several locations throughout the eastern United States from June to August 2019 (Table S4 and Fig. S2). In both species, females tend to mate once and then lay a single clutch of eggs in one pine branch terminus. Thus, distinct larval clusters typically represent full-sib families. To maintain broad-scale geographic differences in behavior or morphology, we grouped field-collected larvae by state (Michigan, Kentucky, Indiana, and North Carolina) and then reared larvae to adults for mating assays and line propagation. Methods for line propagation are described elsewhere (Harper et al. 2016; Bendall et al. 2017). To minimize evolutionary change in the lab, we used adults that were reared either from wild-caught colonies or first-generation lab colonies.

No-choice mating assays were performed from September to October 2019. We used no-choice assays because they are consistent with mating behaviors in the wild (Benjamin 1955; Wilson et al. 1992). Because reproductive isolation can vary across geographic space (Jiang et al. 2013; Rougemont et al. 2015), we assayed mating outcomes in three different crosses, each containing a different combination of *N. lecontei* and *N. pinetum* adults from different U.S. states: Cross 1 = North Carolina *N. lecontei* x Indiana *N. pinetum*; Cross 2 = Kentucky *N. lecontei* x Kentucky *N. pinetum*; and Cross 3 = Kentucky *N. lecontei* x Michigan *N. pinetum*. Each assay consisted of two arenas (plain white 8 × 11 pieces of printer paper) that were divided into six equally sized sections (dividing lines drawn with a black Sharpie pen; Fig. S3). Each arena was recorded by either a Logitech Carl Ziess Tessar HD 1080p or Microsoft LifeCam Cinema Model 1393 camera. We placed a small petri dish (5 cm x 1.5 cm) in each section of each arena, within which we placed a single male and female. Each assay (pair of arenas) consisted of three replicates of each of the four possible female-male pair types for a particular cross: (1) *N. lecontei* female x *N. lecontei* male, (2) *N. lecontei* female x *N. pinetum* male, (3) *N. pinetum* female x *N. lecontei* male, and (4) *N. pinetum* female x *N. pinetum* male. Based on availability of males and females from each geographic region, we were able to complete 12 assays for Cross 1 (*N* = 36 replicates per pair type), 5 assays for Cross 2 (*N* = 15 replicates per pair type), and 5 assays for Cross 3 (*N* = 15 replicates per pair type).

During each assay, which lasted 2 hours, all mating events were recorded for each male-female pair. Each pair was assigned an arbitrary identifier and observed blind with respect to pair type. For an interaction to be considered a mating, the pair in question had to be properly aligned and physically connected (Fig. 1C) for at least 1 minute to ensure a sufficiently secure attachment for sperm transfer. At the end of each assay, the overall mating outcome of each pair was recorded as a “1” if the pair mated at least once or as a “0” if the pair never mated. All males and all unmated females were immediately preserved in 100% ethanol. Mated females were allowed to lay eggs for further lab line propagation but were preserved live after egg laying (except for 17 females that we were unable to find). To confirm recorded mating outcomes, we reviewed all videos (again, blind with respect to each pairs’ identity) and scored each pair as described above. While reviewing videos, we also counted the number of mating attempts made by each male. For a behavior to be counted as a “mating attempt,” we required that the male curled his abdomen in a “U-shape” under the female’s abdomen in an attempt to initiate copulation. We then coded male behavior in each pair as a binary trait: “1” if the male made any attempt and “0” if the male made no attempts.

We used our mating assays to quantify the strength of prezygotic isolation between *N. lecontei* and *N. pinetum* both within each of the three crosses and globally (all crosses combined), following Sobel and Chen (2014). This method requires observed and expected mating frequencies for conspecific and heterospecific pairs. To obtain the observed frequency of conspecific matings, we divided the total number of conspecific pairs that mated (either *N. lecontei* female x *N. lecontei* male or *N. pinetum* female x *N. pinetum* male) by the total number of attempted conspecific crosses. To obtain the observed frequency of heterospecific matings, we divided the total number of heterospecific pairs that mated (either *N. lecontei* female x *N. pinetum* male or *N. pinetum* female x *N. lecontei* male) by the total number of attempted heterospecific crosses. For the expected conspecific and heterospecific mating frequencies under random mating, we used 0.5. We then calculated reproductive isolation using the following equation:

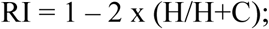

where H = heterospecific pairs and C = conspecific pairs. This equation yields reproductive isolation values ranging from 0 (no reproductive isolation) to 1 (complete reproductive isolation). To further evaluate evidence of prezygotic isolation, we used one-tailed Fisher’s exact tests in each cross and globally (*fisher*.*test* function) to determine whether heterospecific pairs were significantly less likely to mate than conspecific pairs. The Sobel and Chen (2014) equation and our statistical analysis ignores possible asymmetries within pair types. Therefore, to determine whether mating outcomes differed between the four different pair types (*N. lecontei* female x *N. lecontei* male; *N. lecontei* female x *N. pinetum* male; *N. pinetum* female x *N. lecontei* male; *N. pinetum* female x *N. pinetum* male), we also used the *fisher*.*multcomp* function from the RVAideMemoire v0.9-81-2 package, with FDR correction for multiple testing. For plotting proportion data, we used the DescTools v0.99.45 package to calculate 95% Clopper-Pearson confidence intervals for each cross and pair type.

To determine whether body size influenced mating outcomes, we measured body size for *N* = 511 preserved males and females from our mating assays. Because body length is likely to be especially important for proper alignment during mating (Fig. 1C and Video S1), we used body length as our measure of body size. Specifically, we used a Neiko Tools Digital Caliper (model 01407A) to measure each individual from the tip of the head to the tip of the abdomen. To reduce measurement error, each sawfly was measured independently by two individuals and the average body length (in mm) was used. To calculate a size differential for each pair, we subtracted the male body length from the female body length.

We then used logistic regression to evaluate how species, pair type, and size differences affect binary mating outcomes in each of the three crosses and globally. For each cross, we modeled mating outcome as a function of female species, male species, female species x male species interaction, and size differential. For the global model, cross was included as a random effect. Then, we used Type III ANOVAs to evaluate the significance of model terms. If the two species differ in willingness/motivation to mate, we expected significant male species or female species terms. If the type of male-female pair (conspecific versus heterospecific) affects mating outcomes independent of body size, we expected significant female species x male species interaction terms. And if there is size-based assortative mating independent of male-female pair type, we expected significant effects of size differentials in our models, with pairs that mated having a smaller size differential than those that did not. We also used logistic regression to evaluate how species, pair type, and size differences affect male mating attempts (coded as a binary trait) in each cross and globally. Then, we used Type III ANOVAs to evaluate the significance of model terms.

To further explore how variation in body size within and between species relates to variation in the strength of prezygotic isolation among the three crosses, we compared the size differences for all intraspecific pairs (*N. lecontei* female x *N. lecontei* male and *N. pinetum* female x *N. pinetum* male) to the size differences for all interspecific pairs (*N. lecontei* female x *N. pinetum* male and *N. pinetum* female x *N. lecontei* male) within each cross and globally. We then performed one-sided t-tests using the *t*.*test* function to evaluate whether the size differential for interspecific pairs was significantly greater than the size differential for intraspecific pairs.

## Results

### Geographic variation in host needle width and adult body size

Range-wide, needle widths differed significantly among pine species (*P* = 0.000057) and although there was not a general latitudinal effect across all hosts (*P* = 0.34), there was a significant species x latitude interaction (*P* = 0.0000084), indicative of species-specific geographic clines in needle width (Table S5). Post-hoc tests comparing needle width between all pairs of pine species revealed that *P. strobus* (*N. pinetum* host) differed significantly from all *Pinus* hosts associated with *N. lecontei*, except *P. clausa* (sand pine), a thin-needled southern pine that is outside of the range of *N. pinetum* (Table S6) and not considered a preferred/primary host for *N. lecontei* (Wilson et al. 1992; Linnen and Farrell 2010). Ignoring pine species, when we grouped pines by sawfly species instead, we found a significant geographic cline in needle width for pines used by *N. lecontei* (*R*^*2*^ *=* 0.24; *P <* 2.2×10^−16^), but not for the pine species used by *N. pinetum* (*R*^*2*^ *=* 0.001; *P =* 0.78). Overall, needle widths increased with latitude for *N. lecontei*-associated pines, but not *N. pinetum*-associated pines (Fig. 2A). We caution, however, that limited geographical sampling of *P. strobus* needles, especially in northern locales, may have precluded us from detecting geographic clines in needle width.

**Figure 2.**
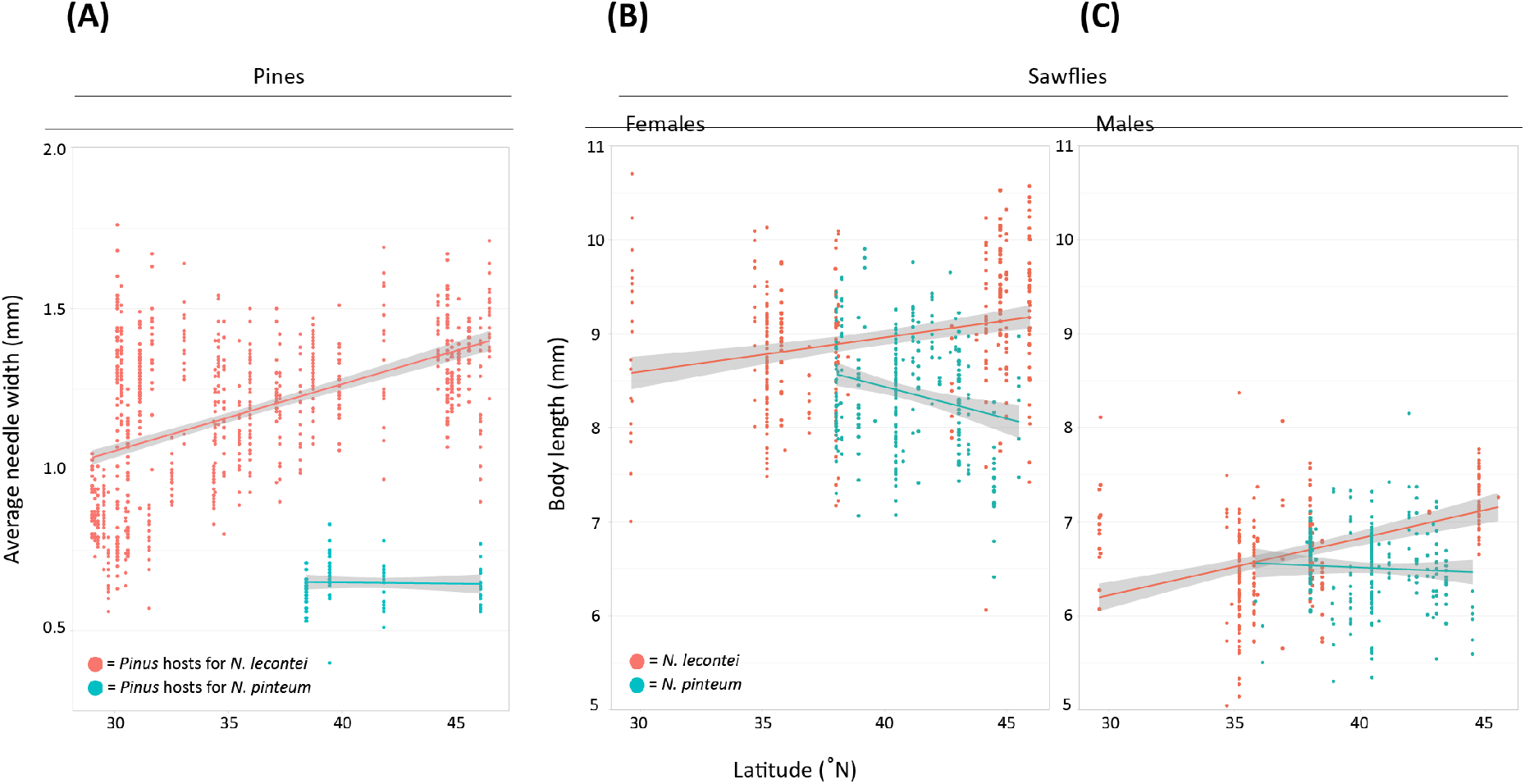
Geographic variation in pine needle width and adult sawfly body size. (A) Clinal variation in needle width (in mm) for 10 *Pinus* species used as hosts by *Neodiprion lecontei* (*P. taeda, P. palustris, P. echinata, P. elliottii, P. clausa, P. glabra, P. virginiana, P. rigida, P. resinosa*, and *P. banksiana*; coral) and for *P. strobus*, the only host used by *N. pinetum* (teal). (B) Clinal variation in body length for *N. lecontei* and *N. pinetum* females. (C) Clinal variation in body length for *N. lecontei* and *N. pinetum* males. In all panels, lines are linear regression models, with gray shading depicting the 95% confidence intervals on the fitted values.

For adult sawfly body size, all model terms and interactions were significant (Table S7), indicating that body size depends on sex, species, and latitude, and that sexes and species both respond differently to latitude. Independent of latitude, *N. lecontei* adults tended to be larger than *N. pinetum* adults and females tended to be larger than males (Fig. 2B, 2C). For *N. lecontei*, we found significant geographic clines in body size for both females (*R*^*2*^*=*0.074; *P =* 5.7 × 10^−7^) and males (*R*^*2*^*=*0.16; *P =* 6.5 × 10^−11^). As was observed for pine needle width, *N. lecontei* size increased with latitude for adult females (Fig. 2B) and adult males (Fig. 2C). In addition, estimated slopes were similar for clinal models of needle width (*m* = 0.021), adult *N. lecontei* females (*m* = 0.039), and adult *N. lecontei* males (*m* = 0.058) (Fig. 2). For *N. pinetum*, there was a significant geographic cline for females (*R*^*2*^ *=* 0.050; *P =* 0.00017), but not males (*R*^*2*^ *=* 0.0023; *P =* 0.47). Unlike *N. lecontei* and unlike the primary host pine for *N. pinetum*, the size of *N. pinetum* females decreased from South to North (Fig. 2B). Like the host pine for *N. pinetum* (Fig. 2A), *N. pinetum* males did not exhibit clinal variation in body size (Fig. 2C).

### Divergent selection on egg size

Egg size differed significantly between *N. lecontei* and *N. pinetum* (linear mixed model, species: χ^2^ = 4.60, *P* = 0.032). As expected from having wider needles and larger adults (Fig. 2), *N. lecontei* eggs were larger than *N. pinetum* eggs (Fig. 3C). Analysis of hatching success for eggs laid by *lecontei*-backcross females revealed a significant egg size effect (*P* = 0.0000095), host effect (*P* = 0.0039), and host x egg size interaction (*P =* 0.00066) (Table S8). Plotting the data revealed that hatching success was higher overall for larger eggs and on the thicker-needled jack pine (*P. banksiana*) (Fig. 3D). Consistent with the trade-off hypothesis, egg size had opposing effects on hatching success on the two pine hosts: having larger eggs increased hatching success on *P. banksiana* but decreased hatching success on the thinner-needled *P. strobus* (Fig. 3D).

**Figure 3.**
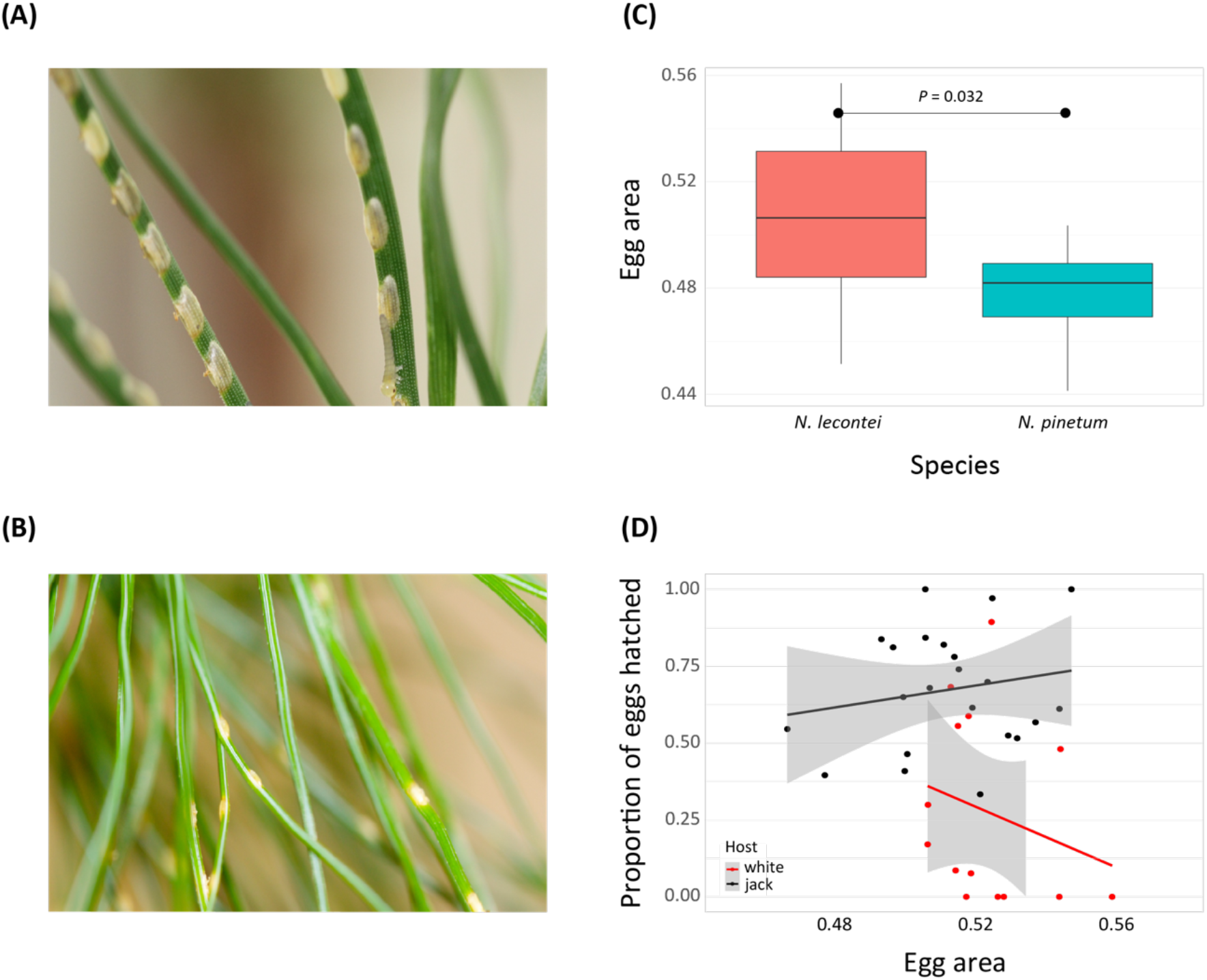
Evidence that egg size is under divergent selection between thick- and thin-needled pines. (A) *Neodiprion lecontei* eggs inside a thicker-needled *Pinus* host (photo by R.K. Bagley). (B) *N. pinetum* eggs inside a thin-needled *Pinus strobus* host (photo by K. Vertacnik). (C) Differences in egg area between *N. lecontei* and *N. pinetum*. Boxes represent interquartile ranges (median ± 2 SD). (D) Hatching success plotted as a function of egg area measured in *N* = 36 recombinant backcross females (individual points). Points are colored depending on whether the female laid her eggs on *P. strobus* (red) or *P. banksiana* (black). Lines are linear regressions to show trends for each host type, with gray shading depicting the 95% confidence intervals on the fitted values. Overall, female egg-laying success increases with egg size on the thicker-needled host (*P. banksiana*), while females that laid bigger eggs had reduced hatching success on the thinner-needled host (*P. strobus*) (see Table S8).

### Size-assortative mating and reproductive isolation

Across all pair types and crosses, mating occurred in 62 of 264 male-female pairs (23.5%). The estimated strength of prezygotic isolation varied across the three different crosses (Fig. 4): prezygotic isolation was strong and significant in Cross 3 (Kentucky *N. lecontei* x Michigan *N. pinetum*; RI = 0.77; *P* = 0.00019), moderate and significant in Cross 2 (sympatric Kentucky *N. lecontei* and *N. pinetum*; RI = 0.37; *P* = 0.047), and weak and not significant in Cross 1 (North Carolina *N. lecontei* x Indiana *N. pinetum*; RI = 0.19; *P* = 0.26). Pairwise comparisons of different pair types within crosses painted a slightly different picture, likely due to asymmetries within conspecific and heterospecific pair types (Table S9 and Fig. 4). For Cross 2, heterospecifics were less likely to mate than conspecifics overall, but none of the pairwise comparisons among pair types were significant after correction for multiple testing (Table S9 and Fig. 4). For Cross 1, which had the largest sample size of any of the crosses, we found significant differences between the two types of conspecific pairs (*N. lecontei* female x *N. lecontei* male versus *N. pinetum* female x *N. pinetum* male, *P* = 0.036) and the two types of heterospecific pairs (*N. lecontei* female x *N. pinetum* male versus *N. pinetum* female x *N. lecontei* male, *P* = 0.0012). In this cross, we also found evidence of asymmetric reproductive isolation: the proportion of pairs that mated differed significantly between conspecific *N. lecontei* pairs and one direction of the interspecific cross (versus *N. lecontei* female x *N. pinetum* male, *P* = 0.00098), but not the other direction (versus *N. pinetum* female x *N. lecontei* male, *P* = 1). Finally, for Cross 3, we found that the proportion of pairs that mated differed between both heterospecific pairs and conspecific pairs involving *N. lecontei* only, but not among any of the other pair types (Fig. 4 and Table S9). Globally, prezygotic isolation was moderate and significant (RI = 0.39; *P* = 0.00038), and all pair type comparisons except *N. pinetum* female x *N. lecontei* male versus *N. pinetum* female x *N. pinetum* male differed significantly in the proportion of pairs that mated (Table S9 and Fig. S4). Overall, these data revealed geographically variable (and sometimes asymmetric) reproductive isolation.

**Figure 4.**
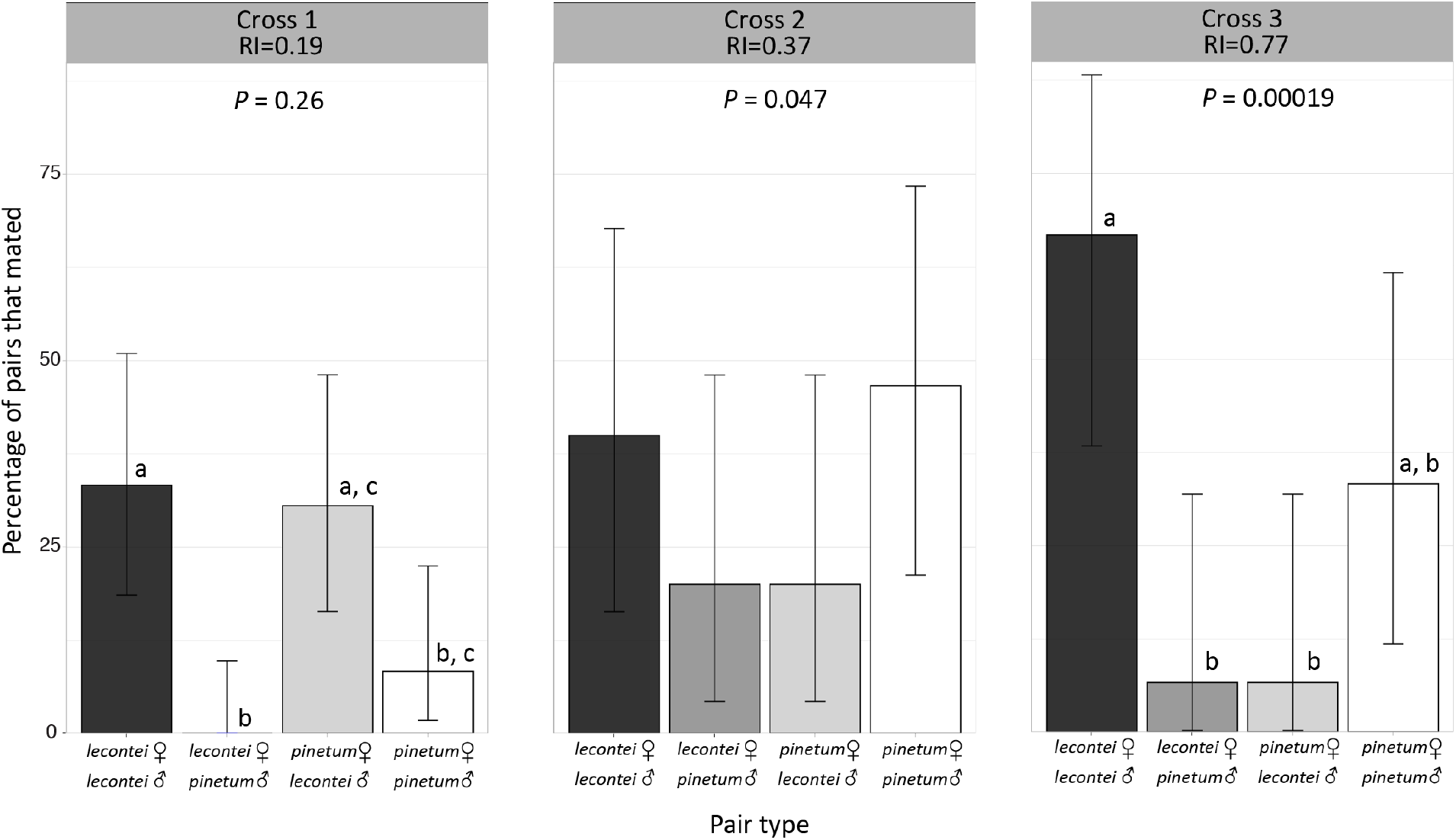
Variation in the strength of prezygotic isolation among three crosses. Each panel gives the percentage of pairs that mated as a function of pair type for one of the three crosses: Cross 1 is North Carolina *Neodiprion lecontei* x Indiana *N. pinetum*; Cross 2 is Kentucky *N. lecontei* x Kentucky *N. pinetum*; Cross 3 is Kentucky *N. lecontei* x Michigan *N. pinetum*. No *N. lecontei* female x *N. pinetum* male pairs mated in Cross 1. Reproductive isolation (RI) values represent the strength of reproductive isolation between *N. lecontei* and *N. pinetum* for each cross as calculated after Sobel and Chen (2014). *P*-values listed for each cross indicates the results of one-sided Fisher’s exact tests; a significant *P*-value indicates that conspecific pairs mated more often than heterospecific pairs. Error bars represent the 95% Clopper-Pearson confidence intervals for the percentage of pairs that mated. Letters indicate pair types that differ significantly at *P* < 0.05 within each cross, after correction for multiple tests (Table S9). Shared letters indicate pair types that were not significantly different. None of the pair types differed significantly in Cross 2.

When we modeled mating outcomes as a function of female species, male species, female species x male species interactions, and size differential, we found that differences in female and male body lengths had a significant effect on mating outcomes in all crosses (Table S10). Plotting the data revealed that, regardless of cross and pair type, pairs that mated were closer in body length (smaller size differentials) than pairs that did not mate (Fig. 5, S5). After accounting for body size differences, we did not detect an additional effect of pair type in any of our crosses, indicating that much of the observed variation in mating outcomes among pair types (and therefore, much of the observed prezygotic isolation) was attributable to body size differences (Table S10). Although size differences appeared to be the most important predictor of mating outcomes in all crosses, we also detected a significant male species effect in Cross 1 (but not the other crosses) and globally (Table S10). This suggests that the observed asymmetries in the proportion of pairs that mated in Cross 1 (Fig. 4) and globally (Fig. S4) may be due to differences in male behavior or morphology. Specifically, pairs that include *N. lecontei* males mated more often than those involving *N. pinetum* males (Fig. 4). When we used male mating attempt rather than mating outcome as our response variable, we found that male species had a significant effect on male mating attempts in Cross 1 and globally (Table S11), and that size differential had a significant effect on male mating attempts in our global analysis only (Table S11). Plotting the data revealed that, in Cross 1 and globally, *N. lecontei* males were more likely to attempt mating than *N. pinetum* males, regardless of female species (Fig. S6). Additionally, regardless of pair type, males were more likely to attempt a mating when size differentials were smaller (Fig. S7), but this effect was modest and only significant in the global analysis (Table S11).

**Figure 5.**
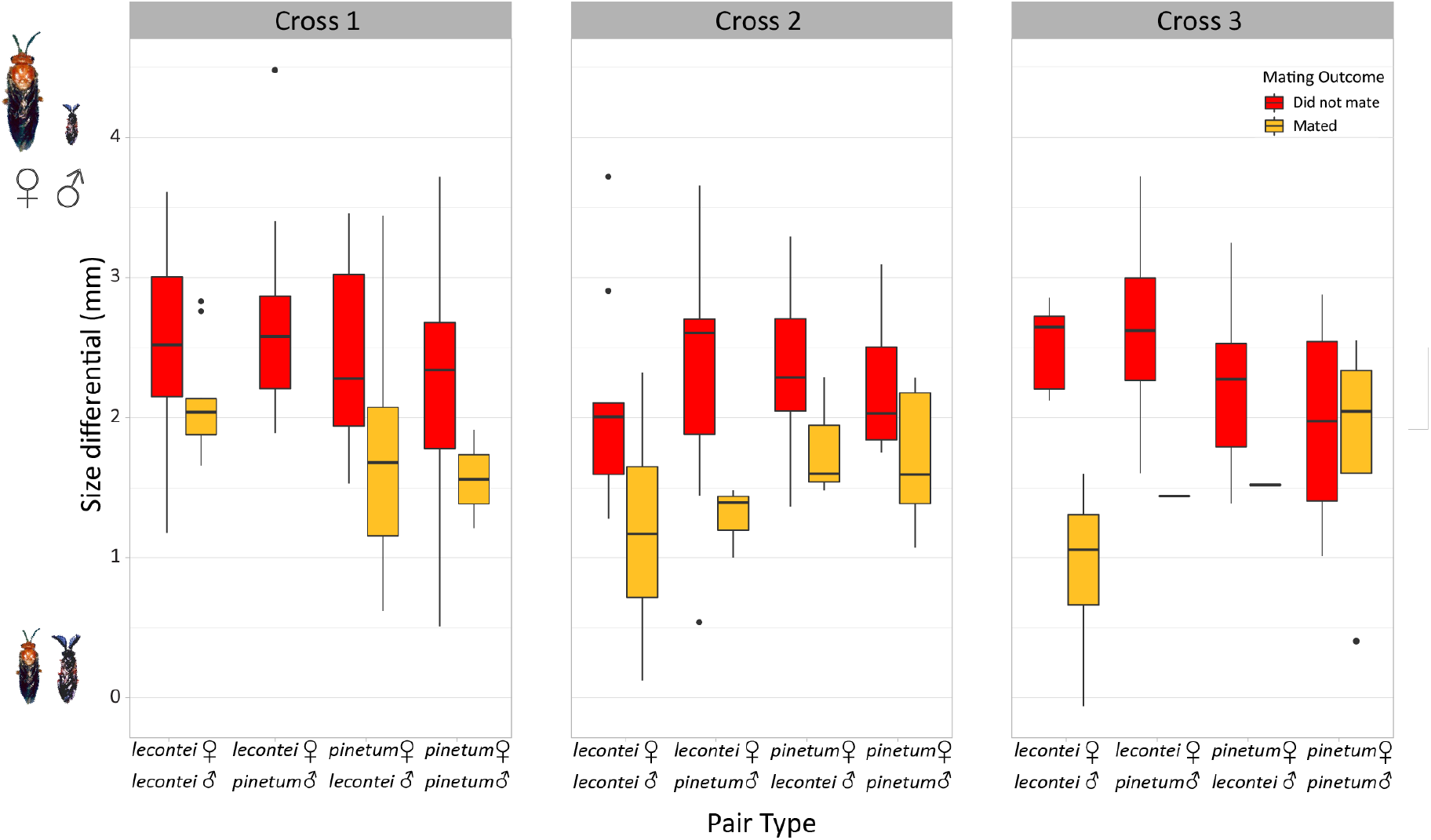
Across all crosses and pair types, pairs that did not mate (red boxes) had greater size differentials than pairs that did mate (yellow boxes). Cross 1 is North Carolina *Neodiprion lecontei* x Indiana *N. pinetum*; Cross 2 is Kentucky *N. lecontei* x Kentucky *N. pinetum*; Cross 3 is Kentucky *N. lecontei* x Michigan *N. pinetum*. No *N. lecontei* female x *N. pinetum* male pairs mated in Cross 1 (no yellow bar). Size differentials are female body length minus male body length; higher values reflect larger size differences (adults not drawn to scale). Boxes represent interquartile ranges (median ± 2 SD), with size differential outliers indicated as points. Note that while we are showing size differentials on the y-axis to visualize the consistency of this pattern, mating outcome was modeled as the response variable. Size differential had a significant effect on mating outcome in all crosses (Table S10).

Variation in body size within and between species may explain variation in the strength of prezygotic isolation among the three crosses. For example, the cross with the highest prezygotic isolation (Cross 3) also had the most pronounced differences between inter- and intra-specific pair types in terms of size differentials (Fig. S8 and Table S12). Conversely, the cross for which we did not detect significant prezygotic isolation (Cross 1) also had the largest *intra*specific size differentials, and interspecific size differentials were not significantly higher than intraspecific size differentials (Fig. S8 and Table S12).

## Discussion

Magic traits are a potentially important mechanism for facilitating speciation-with-gene-flow (Servedio et al. 2011). Body size, in particular, appears to be a widespread magic trait in diverse animal taxa but is relatively unexplored in plant-feeding insects (but see Augustyn et al. 2017), which include some of the most speciose lineages on earth (Misof et al. 2014; Forbes et al. 2018). Here, we demonstrate that two key requirements for body size to be considered a magic trait are met in a pair of recently diverged sawfly species adapted to different pine hosts. First, we provide two lines of evidence that body size is under divergent selection stemming from differences in pine use: (1) body size clines in sawfly adults appear to track clines in needle width for their primary pine hosts (Fig. 2); and (2) performance data from recombinant backcross females suggest that egg size is under divergent selection between thick-needled and thin-needled pines (Fig. 3). Second, our mating assays provide clear evidence that body size differences between the species contribute to prezygotic isolation (Figs. 4, 5). Nevertheless, while our data satisfy several key requirements for body size to act as a magic trait in this system, more work is needed to clarify genetic and ecological mechanisms. Here, we discuss limitations of our current data, as well as broader implications for the role of body size in plant-feeding insect speciation.

### Host-associated divergent selection on body size

Geographic variation in body size is widespread (and variable) in nature (Ashton 2002; Chown and Gaston 2010). In *N. lecontei*, we found that body size increases with latitude in both adult males and females (Fig. 2B, 2C). This pattern is consistent with Bergmann’s rule, which has been documented in diverse animal taxa (Bergmann 1847; Shelomi 2012; Auteri 2022). Although this geographic pattern is widespread, underlying mechanisms appear to vary across taxa (Steudel et al. 1994; Terribile et al. 2009; Chown and Gaston 2010; Stillwell 2010). Because it covaries with latitude, temperature is one such mechanism that may give rise to an increase in body size with latitude. Specifically, because surface-to-volume ratios decline with body size, larger animals may be better able to minimize heat loss in cold environments (Bergmann 1847). But a corresponding latitudinal increase in needle widths for *N. lecontei*’s primary pine hosts suggests an alternative explanation: body size clines in *N. lecontei* are driven by selection to fit within host needles. Clinal variation in host size and constraints imposed by development within host tissue have also been invoked to explain clinal variation in body size in seed beetles (Stillwell et al. 2007) and in camellia weevils (Toju and Sota 2006).

In comparison with *N. lecontei, N. pinetum* exhibits strikingly different adult body size clines. Both females (Fig. 2B) and males (Fig. 2C) tend to become smaller at higher latitudes, a pattern consistent with converse Bergmann’s rule (Mousseau 1997; Ashton and Feldman 2003; Blanckenhorn and Demont 2004; Stillwell et al. 2007), although the latitudinal cline was only significant in females. The typical explanation for the converse Bergmann’s rule is that shorter growing seasons at higher latitudes constrain body size via limited time for growth and reproduction (Blanckenhorn and Demont 2004). As *N. lecontei* and *N. pinetum* are recently diverged sister species (Herrig et al. in prep; Bendall et al. 2022) with very similar life histories, a similar geographic range (Fig. 1A), and overlapping phenologies in most of their range (including northern latitudes; Wilson 1977), it seems unlikely that differences in clinal patterns are due to different plastic or adaptive responses to shared abiotic factors. Instead, the most obvious difference between the two species is that *N. pinetum* specializes on a single pine species that does not appear to exhibit clinal variation in needle width, whereas needle width increases with latitude for *N. lecontei* hosts (Fig. 2A).

Overall, based on the biology and ecology of these species, we hypothesize that the observed geographic clines in body size are primarily due to host-associated divergent selection on body size traits both among *N. lecontei* populations and between *N. lecontei* and *N. pinetum*. However, because our samples were primarily adults that were reared in the lab from field-caught larvae, one non-mutually exclusive explanation for the observed patterns is that the observed clines are due to plastic responses to host plants or the local climate (Davidowitz et al. 2004; Du et al. 2021). Although additional experiments are needed to measure heritability and characterize plasticity for body size traits, our experience to date suggests both that body size traits are highly heritable and that they can be influenced by development on a non-preferred host (unpublished data). Our clinal data are also insufficient for inferring causal mechanisms generating divergent selection on body size; for this, experiments are needed. We consider three non-mutually exclusive mechanisms for divergent selection on body size in sawflies and other plant-feeding insects: trade-offs involving the fit of immature stages within host tissues, trade-offs involving the fit of female ovipositors within hosts, and trade-offs involving visual appearance and movement on the host.

On one side of all potential host-related body size trade-offs are the myriad of abiotic and biotic selective pressures that favor large body sizes in egg, juvenile, or adult life stages. For example, larger eggs are likely to provide more resources for an embryo to complete development, hatch, and for the newly hatched larva to migrate to a suitable feeding location on the host (Davis et al. 2012). Consistent with this prediction, we found that when females laid eggs on the thicker-needled pine, the proportion of eggs that hatched and survived the first hours of larval life (up to 48 hours) increased with egg size (Fig. 3D). Having a head start on growth may also enable individuals to attain a larger adult size when growing time is limited (Azevedo et al. 1996; Meister et al. 2018). Larger females may, in turn, produce larger or more eggs (Pincheira-Donoso and Tregenza 2011; Davis et al. 2012). Positive relationships between female body size and fecundity are well documented in diverse insect taxa (Honěk 1993), including *Neodiprion lecontei* (Harper et al. 2016). Larger males may be favored via sexual selection (Chown and Gaston 2010). Finally, larger body size in life stages that overwinter may increase survival rates (Piiroinen et al. 2011; Kovacs and Goodisman 2012).

Although many selective mechanisms favor increases in body size, there are also constraints. For example, many different types of insects develop partly or wholly within the tissue of their living host plant or, analogously, host animal (e.g., parasitoids). Whenever this occurs, host size will place substantial constraints on egg size and body size in immature stages (Hardy et al. 1992; Mackauer and Chau 2001; Stillwell et al. 2007; Chown and Gaston 2010), which may set an upper limit on adult size (Meister et al. 2018). Consistent with this expectation, our backcross data show that females that produce bigger eggs have reduced hatching success on a thin-needled pine (Fig. 3D). While we have not experimentally tested the precise mechanism of hatching failure, observations of successful egg development in pine needles provide some clues. Normally, eggs are not visible in egg pockets (i.e., the niches carved into the needles by the female, Fig. 1B) until right before hatching (∼2 weeks post-oviposition), when the eggs swell and cause the egg pocket to split open. Two potential size-related causes for hatching failure on thin needles are: (1) large eggs do not fit entirely within egg pockets (see Fig. 1B), and air exposure causes eggs to desiccate and die; or (2) large eggs cause egg pockets to gape open too early, causing the egg-bearing needle to dry out before embryonic development is complete. Careful observation of egg development within needles and experimental manipulation of egg pockets or host needles are needed to evaluate these possibilities.

A good match between adult female ovipositor size and host size is also essential to successful reproduction in many plant-feeding insects (Sota et al. 2007; Holma 2009) and parasitoids (Luz et al. 2020), and thin, delicate host tissues may place an upper limit on ovipositor size. In support of this prediction, *Plateumaris constricticollis toyamensis* leaf beetles that lay their eggs in thin-stemmed hosts have shorter ovipositors than a closely related subspecies (*P. c. babai*) that lays its eggs in hosts with thicker stems (Sota et al. 2007). This pattern is also observed in *Neodiprion*. In a previously published experiment involving *lecontei-* backcross females, it was demonstrated that females with larger ovipositors had reduced oviposition success on the thin-needled white pine host of *N. pinetum* (Bendall et al. 2017). Observations of egg-bearing needles suggest that the mechanism responsible is that large ovipositors cause the female to cut through the host tissue, leading to needle desiccation and egg death. Because ovipositor size is often directly related to adult female body size (Sota et al. 2007; Yanagi and Tuda 2012), constraints on ovipositor size are also likely to constrain adult female body size.

Finally, a third potential source of host-associated divergent selection on body size is via how it affects visual contrast or movement on the host plant. In cryptic species, individuals with body sizes that are visually mismatched with their host background may experience increased predation (Sandoval 1994a, b; Nosil 2007; Sandoval and Crespi 2008). For example, great tits (*Parus major*) preferentially attack larger cryptic models of lepidopteran larvae (Mänd et al. 2007). Similarly, cryptic *Cephalelus uncinatus* leafhoppers avoid avian predation by resting on host plants with stems that match their body size: larger individuals are always found on plants with thicker stems, while smaller individuals are found on plants with thinner stems (Augustyn et al. 2017). By contrast, in aposematic species such as *N. lecontei* and *N. pinetum* (Lindstedt et al. 2022), having a larger visual signal (either via larger bodies or larger groups) may enhance predator avoidance learning (Mappes and Alatalo 1997; Gamberale and Tullberg 1998; Forsman and Merilaita 1999; Riipi et al. 2001). However, group or individual size may be limited by physical properties of the host. White pine needles are much thinner and more flexible than those of many other pines, which may limit the number or size of larvae that can feed in clusters. Body size can also affect how well insects move on the host. For example, experimental evolution of pigeon lice on differently sized pigeons produced rapid divergence in body size. The proposed selective mechanism: on small pigeon hosts, small lice are better able to avoid death by preening because they can hide between feather barbs; on large pigeon hosts, larger lice are both more fecund and better able to avoid preening because they move more quickly between feather barbs (Villa et al. 2019). Overall, our work (together with an extensive literature on the ecology of insect body size) highlights the potential for body size to be a near-ubiquitous target of divergent selection when insect populations adapt to different host plants.

### Size-assortative mating and reproductive isolation

Size-assortative mating within and between species is common in a variety of taxa (e.g., Harari et al. 1999, Jones et al. 2003, McKinnon et al. 2004, Bearhop et al. 2005, Rougemont et al. 2015, Greenway et al. 2016). Consistent with this body of work, we found that, regardless of female-male pair type, pairs that were more dissimilar in size were less likely to mate (Fig. 5). Size differences also appear to contribute to significant, but asymmetric and geographically variable reproductive isolation between *N. pinetum* and *N. lecontei* that is maximized: (1) in the direction of the cross that is most dissimilar in size (*N. lecontei* females and *N. pinetum* males; Fig. 4) and (2) in crosses where size differences between species are greater than those within species (Fig. S8). These findings predict that introgression in nature is likely to be asymmetric, a prediction that is supported by estimated gene flow rates from sympatric populations of *N. lecontei* and *N. pinetum* from Kentucky (Bendall et al. 2022). Combined with clinal patterns in adult body size (Fig. 2B, 2C), our findings also predict that gene flow in nature is likely to be lowest in northern parts of the shared range, where size differences between the species are greatest. This prediction can be tested via demographic analysis of northern populations.

At a mechanistic level, Crespi (1989) proposed three hypotheses that can explain size-assortative mating in insects: (1) mate availability; (2) mating constraints; and (3) mate choice. Mate availability generates size-assortative mating when individuals group together by size or when female and male body size covaries in time (Miyashita 1994; Harari et al. 1999). Mating constraints generate size-assortative mating when mating is physically difficult. This can be due to poor alignment during mating due to mismatched female and male body sizes (e.g., Weissman et al. 2008; Villa et al. 2019) and/or due to mismatched female and male genitalia (e.g., Tanabe and Sota 2008; Masly 2012). Mate choice can generate size-assortative mating if either or both sexes select large mates (e.g., Rowe and Arnqvist 1996) and, consequently, male-male competition allows larger males to win access to the large females (e.g., Harari et al. 1999; Villa et al. 2019).

Our data indicate that in no-choice mating assays, mating outcomes in both conspecific and heterospecific pairs of *Neodiprion* sawflies are strongly dependent on how much bigger the female is than the male. Based on observing mating interactions, this size-assortative mating appears to be due, at least in part, to difficulty of obtaining the correct body alignment when female and male body sizes are mismatched. Additionally, females frequently resist male mating attempts and can physically displace comparatively small males that are attempting to mate, sometimes even attacking the males and tearing off antennae or legs (Glover, personal observation; Wilson et al. 1992). However, when the male body size is more closely matched to that of the female, the male can withstand female resistance and properly connect his lower abdomen with the female’s lower abdomen (Video S1).

Mate choice may also contribute to size-assortative mating. For example, in the field, males often mate with multiple females, while females typically mate with a single male (Wilson et al. 1992). Thus, choosy males that avoid attempting to mate with comparatively large females may have a fitness advantage because they are less likely to be maimed or killed, thereby preventing them from pursuing additional females. Consistent with this prediction, in at least one of our crosses (Cross 1), there was a significant effect of male species on mating outcome (Table S10) and male mating attempts (Table S11), with the smaller species (*N. pinetum*) seemingly more reluctant to mate overall (Figs. 4, S6). However, there was only a significant effect of size differential on male attempts in our global analysis (Table S11), which suggests that mating outcomes are not primarily driven by male choice. Although we did not test for size-assortative mating due to mate availability or genital mismatch, it is possible that these mechanisms are also contributing to mating outcomes between *N. lecontei* and *N. pinetum*.

Interestingly, while our model results suggest that the specific pairing of male species and female species did not affect mating outcomes once size was accounted for (Table S10), other patterns in our data hint that body size is not the full story. For example, in the two crosses with stronger prezygotic isolation (Cross 2 and Cross 3), both interspecific pair types mate at similar frequencies despite asymmetric interspecific size differentials (Figs. 4, 5). Specifically, because *N. lecontei* are generally larger than *N. pinetum*, pairs consisting of *N. lecontei* females and *N. pinetum* males tend to have larger size differentials than pairs consisting of *N. pinetum* females and *N. lecontei* males. Therefore, if size differential is the only factor determining mating outcome, *N. pinetum* female x *N. lecontei* male pairs (smaller size differential) should mate more readily than *N. lecontei* female x *N. pinetum* male pairs (larger size differential). Yet we only observe this predicted asymmetry in Cross 1, which is also the only cross involving allopatric populations of both species (note that while populations for Cross 3 were sampled from different areas, both populations co-occur with heterospecific populations). These patterns are consistent with asymmetric reinforcement (Yukilevich 2012; Greenway et al. 2016): to avoid maladaptive hybridization (Bendall et al. 2017), *N. pinetum* females that encounter *N. lecontei* males may have evolved increased mate discrimination, likely via some mechanism other than body size.

It is also possible that the artificial mating assay conditions either exaggerated the effect of body size or minimized the effects of other chemical, behavioral, and morphological traits that affect mating outcomes in nature (Coyne et al. 2005; Xue et al. 2014; Xu and Turlings 2018). Thus, more realistic mating assays (e.g., in the presence of the host, under field conditions) would be a fruitful avenue for future work. Additionally, our assays do not preclude the existence of other mechanisms of prezygotic isolation (and other magic traits) in nature. Although female pheromone composition and male response to female pheromones appear to be similar between *N. lecontei* and *N. pinetum* (Matsumura et al. 1979; Kraemer et al. 1979, 1981, 1984; Anderbrant 1993), there could be assortative mating by preferred host. Like many other insects, *Neodiprion* adults tend to mate on the host plant; thus, changes in host preference could produce non-random mating (e.g., as in *Rhagoletis pomonella;* Feder et al. 1994; Feder 1998; Matsubayashi et al. 2010). That said, although adult females exhibit strong host preferences (Bendall et al. 2017), field and behavioral data suggest that adult males may not rely on host cues to locate females (Linnen, unpublished data). Since male behavior determines heterospecific encounter rates in *Neodiprion*, divergent host preferences may not produce assortative mating, although further work is needed to confirm this. Differences in phenology (e.g., if males have different emergence peaks or flight activity patterns), which may or may not be host related, could also result in non-random mating and temporal isolation (Zhang et al. 2019; Larson et al. 2019). Thus, more work is needed to evaluate the importance of size relative to other potential sources of non-random mating in *Neodiprion* sawflies.

### Is body size really “magic” in pine sawflies?

Although “magic trait” studies are generally studied at the phenotypic level, the “magic” that facilitates speciation-with-gene-flow is pleiotropy: the trait under divergent selection and the trait causing non-random mating must share a common genetic basis (Servedio et al. 2011). While our data support several magic trait predictions, we have not experimentally manipulated traits or provided direct evidence of pleiotropy in this study. For example, while data in many arthropods support our assumption that larger eggs produce larger larvae and larger adults (Fox and Czesak 2000; Macke et al. 2011), we have not confirmed this in *Neodiprion*. One strategy for doing so would be to track individual eggs from hatching to adulthood, a difficult experiment in these highly gregarious species that do not fare well when isolated. An alternative approach would be to use quantitative genetic approaches to evaluate genetic correlations between egg size and adult body size or colocalization of quantitative trait loci for egg size and adult body size. There is, however, already some evidence that adult female body size and ovipositor size are genetically correlated (via pleiotropy or tight linkage): the two traits are positively correlated in recombinant backcross females (Bendall et al. 2017). Overall, much work remains to determine the strength and underlying causes of genetic correlations among body size traits expressed at different developmental stages (eggs, larvae, and adults) and between the sexes. Additional work is also needed to evaluate the “effect size” (i.e., how much an allele or trait increases reproductive isolation) of body size relative to other reproductive barriers and other potential magic traits (Servedio et al. 2011; Nosil and Schluter 2011). Fortunately, growing genomic resources (Vertacnik et al 2017; Herrig et al. 2021) and the ability to rear and cross these species in the lab (Bendall et al. 2017) make this a highly tractable system for uncovering the genetic and ecological mechanisms underlying magic traits and speciation.

## Conclusions

A vast literature documents the importance of body size to survival and reproductive success in diverse taxa through a variety of mechanisms (Honěk 1993; Davidowitz et al. 2004; Chown and Gaston 2010; Seifert et al. 2022). In plant-feeding insects (and many other parasitic organisms that specialize on particular hosts) the need to fit entire bodies or body parts (e.g., ovipositors) within the host tissue places a particularly strong constraint on body size. We argue that this constraint, likely unique to parasitic lifestyles, may make body size a common magic trait in plant-feeding insects and other host-specialized parasites. In support of this hypothesis, we provide several lines of evidence that suggest body size is a magic trait in pine sawflies. However, much work remains to clarify the role of pleiotropy and the importance of body size relative to other reproductive barriers in this system, and these studies need to be replicated in diverse taxa to establish the general importance of body size as a speciation trait. Together, such studies would provide great insight into why plant-feeders and parasitoids are so unusually diverse (Forbes et al. 2017). Additionally, our work highlights the importance of studying spatial variation in morphology, ecology, and reproductive isolation. Understanding how and why patterns of reproductive isolation vary across geographic space can not only provide important insights into the nature of species boundaries but can also allow us to predict future hybridization dynamics. This information has basic and applied research value (e.g., hybridizing insect pest management).

## Author Contributions

ANG and CRL participated in the design of the study, data analysis and drafting of the manuscript. ANG, AW, AF, EEB, JWT2 and GN participated in data collection. All authors have read and approved the final manuscript.

## Acknowledgments

We would like to thank Dr. Robin Bagley, Allyson Appel, and Maya Woolfolk for assistance with specimen collection. We thank Alexandria Pete and Sarah Lau for assistance with data collection and members of the Linnen lab for insect rearing assistance. We also thank members of the Linnen lab for helpful comments and discussion that improved the quality of this manuscript. This research was supported by the University of Kentucky Merit Fellowship to ANG, University of Kentucky Ribble Travel Grant to JWT2, University of Kentucky Undergraduate Summer Research and Creativity Grant to GN, and the National Science Foundation DEB-1257739 and DEB-CAREER-1750946 to CRL.

## Data Accessibility Statement

All raw data spreadsheets and R code will be archived on Dryad upon publication.

