## Supporting Information for "Body size as a magic trait in two plant-feeding insect species"

**Table S1.** Sampling locations of needles from 11 *Pinus* species

**Table S2.** Sampling locations for all adults measured to characterize geographic variation in body length

**Table S3.** Sampling locations of all *Neodiprion lecontei* and *N. pinetum* females from which eggs were extracted and measured to test for differences in egg size between the species

**Table S4.** Sampling locations of all colonies used in no-choice mating assays

**Table S5.** Type III ANOVA table for linear regression model for pine needle width

**Table S6.** Post-hoc estimated marginal means for comparisons of needle width between all pairs of 11 *Pinus* pine species

**Table S7.** Type III ANOVA table for linear regression model for adult sawfly body size

**Table S8.** Type III ANOVA table for logistic regression model for hatching success modeled as a function of egg area and host

**Table S9.** Variation in the strength of prezygotic isolation among crosses and among different pair types within each cross

**Table S10.** Type III ANOVA tables for logistic regression model for binary mating outcome modeled as a function of female species, male species, female species x male species interaction, and size differential (female body length – male body length)

**Table S11.** Type III ANOVA tables for logistic regression model for binary male mating attempts modeled as a function of female species, male species, female species x male species interaction, and size differential (female body length – male body length)

**Table S12.** Results of one-sided t-tests testing whether heterospecific pairs had significantly greater size differentials than conspecific pairs

**Figure S1.** Map depicting the sampling locations of all adults measured to characterize geographic variation in body length

**Figure S2.** Map depicting the sampling locations of all colonies used in no-choice mating assays

**Figure S3.** Set-up of each arena in no-choice mating assays

**Figure S4.** Prezygotic isolation between *Neodiprion lecontei* and *N. pinetum*

**Figure S5.** Size differentials of pairs that mated compared to pairs that did not mate for each pair type for all crosses combined (globally)

**Figure S6.** Variation in male willingness/motivation to mate

**Figure S7.** Size differentials of pairs where no male attempt to mate was made compared to pairs where an attempt was made for each pair type within each cross and for all crosses combined (globally)

**Figure S8.** Size differentials for interspecific pairs compared to intraspecific pairs for each cross and globally (all crosses combined)

**Video S1.** A *Neodiprion lecontei* mating pair

**Table S1. Sampling locations of trees from which needle width was measured for 11 *Pinus* species: white (*P. strobus*), sand (*P. clausa*), slash (*P. elliottii*), spruce (*P. glabra*), loblolly (*P. taeda*), longleaf (*P. palustris*), shortleaf (*P. echinata*), virginia (*P. virginiana*), pitch (*P. rigida*), red (*P. resinosa*), and jack (*P. banksiana*).**

| Common Name | Species Name | Latitude | Longitude | Collection Date |
|---|---|---|---|---|
| white | *P. strobus* | 38.423927 | -82.323034 | 5/15/17 |
| white | *P. strobus* | 39.458826 | -77.98593 | 5/17/17 |
| white | *P. strobus* | 41.846502 | -70.679302 | 5/18/17 |
| white | *P. strobus* | 46.105318 | -90.987811 | 5/21/17 |
| white | *P. strobus* | 38.423508 | -82.32448 | 8/10/17 |
| white | *P. strobus* | 39.458802 | -77.986075 | 8/11/17 |
| white | *P. strobus* | 41.846617 | -70.679839 | 8/12/17 |
| white | *P. strobus* | 46.105431 | -90.987594 | 8/15/17 |
| sand | *P. clausa* | 29.500249 | -81.847438 | 5/9/17 |
| sand | *P. clausa* | 29.250293 | -81.727206 | 5/9/17 |
| sand | *P. clausa* | 29.119316 | -81.576044 | 5/9/17 |
| sand | *P. clausa* | 29.004558 | -81.757403 | 5/9/17 |
| sand | *P. clausa* | 29.502104 | -81.85023 | 8/4/17 |
| sand | *P. clausa* | 29.25056 | -81.748781 | 8/4/17 |
| sand | *P. clausa* | 29.119401 | -81.576066 | 8/4/17 |
| sand | *P. clausa* | 29.004634 | -81.757364 | 8/4/17 |
| slash | *P. elliottii* | 31.634687 | -83.584268 | 5/8/17 |
| slash | *P. elliottii* | 30.283531 | -82.475735 | 5/8/17 |
| slash | *P. elliottii* | 30.098727 | -83.469845 | 5/9/17 |
| slash | *P. elliottii* | 31.101918 | -86.558239 | 5/10/17 |
| slash | *P. elliottii* | 31.634689 | -83.584249 | 8/3/17 |
| slash | *P. elliottii* | 30.28378 | -82.474261 | 8/3/17 |
| slash | *P. elliottii* | 30.09867 | -83.469834 | 8/5/17 |
| slash | *P. elliottii* | 31.101896 | -86.558168 | 8/5/17 |
| spruce | *P. glabra* | 29.708029 | -82.452411 | 5/9/17 |
| spruce | *P. glabra* | 30.103285 | -83.558538 | 5/9/17 |
| spruce | *P. glabra* | 30.473062 | -84.933068 | 5/10/17 |
| spruce | *P. glabra* | 31.49928 | -84.593243 | 5/10/17 |
| spruce | *P. glabra* | 29.707558 | -82.452137 | 8/5/17 |
| spruce | *P. glabra* | 30.103315 | -83.558482 | 8/5/17 |
| spruce | *P. glabra* | 30.573921 | -84.703946 | 8/5/17 |
| spruce | *P. glabra* | 31.50002 | -84.593168 | 8/5/17 |
| loblolly | *P. taeda* | 34.563097 | -84.944423 | 5/7/17 |
| loblolly | *P. taeda* | 34.412177 | -81.706041 | 5/7/17 |

| | | | | |
|---|---|---|---|---|
| loblolly | *P. taeda* | 30.572694 | -84.704846 | 5/10/17 |
| loblolly | *P. taeda* | 35.941871 | -86.527063 | 5/10/17 |
| loblolly | *P. taeda* | 34.80557 | -77.155363 | 5/16/17 |
| loblolly | *P. taeda* | 34.563783 | -84.944234 | 8/2/17 |
| loblolly | *P. taeda* | 34.412076 | -81.706054 | 8/2/17 |
| loblolly | *P. taeda* | 30.57394 | -84.703948 | 8/5/17 |
| loblolly | *P. taeda* | 35.941979 | -86.527703 | 8/6/17 |
| loblolly | *P. taeda* | 34.805543 | -77.155246 | 8/11/17 |
| longleaf | *P. palustris* | 33.050526 | -83.71877 | 5/8/17 |
| longleaf | *P. palustris* | 30.283531 | -82.475735 | 5/8/17 |
| longleaf | *P. palustris* | 30.103285 | -83.558538 | 5/9/17 |
| longleaf | *P. palustris* | 31.097964 | -86.555047 | 5/10/17 |
| longleaf | *P. palustris* | 33.052152 | -83.71943 | 8/3/17 |
| longleaf | *P. palustris* | 30.28378 | -82.474261 | 8/3/17 |
| longleaf | *P. palustris* | 30.103315 | -83.558482 | 8/5/17 |
| longleaf | *P. palustris* | 31.098276 | -86.556008 | 8/5/17 |
| shortleaf | *P. echinata* | 32.507717 | -83.453204 | 5/8/17 |
| shortleaf | *P. echinata* | 34.35316 | -87.508586 | 5/10/17 |
| shortleaf | *P. echinata* | 35.4746219 | -80.261984 | 5/15/17 |
| shortleaf | *P. echinata* | 32.507293 | -83.453275 | 8/3/17 |
| shortleaf | *P. echinata* | 34.35306 | -87.507954 | 8/6/17 |
| shortleaf | *P. echinata* | 35.474506 | -80.262149 | 8/10/17 |
| virginia | *P. virginiana* | 35.941871 | -86.527063 | 5/10/17 |
| virginia | *P. virginiana* | 38.167388 | -83.590475 | 5/15/17 |
| virginia | *P. virginiana* | 37.11326 | -78.027123 | 5/16/17 |
| virginia | *P. virginiana* | 38.728319 | -79.463185 | 5/17/17 |
| virginia | *P. virginiana* | 35.941979 | -86.527703 | 8/6/17 |
| virginia | *P. virginiana* | 38.16747 | -83.590316 | 8/10/17 |
| virginia | *P. virginiana* | 37.112883 | -78.026644 | 8/11/17 |
| virginia | *P. virginiana* | 38.728206 | -79.463082 | 8/11/17 |
| pitch | *P. rigida* | 38.728319 | -79.463185 | 5/17/17 |
| pitch | *P. rigida* | 39.885551 | -74.506235 | 5/17/17 |
| pitch | *P. rigida* | 41.846502 | -70.679302 | 5/18/17 |
| pitch | *P. rigida* | 37.262241 | -84.96338 | 5/21/17 |
| pitch | *P. rigida* | 37.262241 | -84.96338 | 8/6/17 |
| pitch | *P. rigida* | 38.728206 | -79.463082 | 8/11/17 |
| pitch | *P. rigida* | 39.885802 | -74.505763 | 8/12/17 |
| pitch | *P. rigida* | 41.846617 | -70.679839 | 8/12/17 |
| red | *P. resinosa* | 44.6381 | -84.632676 | 5/20/17 |
| red | *P. resinosa* | 46.095588 | -85392621 | 5/20/17 |

| | | | | |
|---|---|---|---|---|
| red | *P. resinosa* | 44.844026 | -88.449944 | 5/20/17 |
| red | *P. resinosa* | 45.140157 | -91.935225 | 5/21/17 |
| red | *P. resinosa* | 44.637999 | -84.632782 | 8/14/17 |
| red | *P. resinosa* | 46.094974 | -85.391598 | 8/14/17 |
| red | *P. resinosa* | 44.84418 | -88.449958 | 8/14/17 |
| red | *P. resinosa* | 45.140059 | -91.934999 | 8/15/17 |
| jack | *P. banksiana* | 44.6381 | -84.632676 | 5/20/17 |
| jack | *P. banksiana* | 45.602126 | -89.352201 | 5/20/17 |
| jack | *P. banksiana* | 46.500632 | -90.462546 | 5/21/17 |
| jack | *P. banksiana* | 44.226623 | -90.707503 | 5/21/17 |
| jack | *P. banksiana* | 44.637999 | -84.632782 | 8/14/17 |
| jack | *P. banksiana* | 45.602341 | -89.353138 | 8/14/17 |
| jack | *P. banksiana* | 46.500615 | -90.463138 | 8/15/17 |
| jack | *P. banksiana* | 44.226879 | -90.707916 | 8/15/17 |

**Table S2. Sampling locations for all *Neodiprion lecontei* and *N. pinetum* individuals measured to characterize geographic variation in body size.** Individuals used in the mating assays are included. *N* represents the number of individual females and males collected from each sampling site within each region.

| State | Species | Latitude | Longitude | *N* Females | *N* Males |
|---|---|---|---|---|---|
| Alabama | *N. lecontei* | 34.8935 | -86.4459 | 0 | 1 |
| Florida | *N. lecontei* | 29.624 | -82.3082 | 8 | 10 |
| | | 29.6703 | -82.2558 | 4 | 1 |
| | | 29.6694 | -82.2556 | 7 | 3 |
| Indiana | *N. pinetum* | 39.1871 | -86.5447 | 5 | 0 |
| | | 40.4693 | -86.9132 | 73 | 85 |
| | | 41.1655 | -87.2717 | 19 | 9 |
| | | 41.3811 | -85.0525 | 1 | 0 |
| | | 41.3823 | -85.0534 | 8 | 0 |
| Kentucky | *N. lecontei* | 37.9868001 | -84.416332 | 1 | 0 |
| | | 38.0146657 | -84.503891 | 35 | 49 |
| | | 38.09375 | -84.554583 | 1 | 5 |
| | | 38.1074375 | -84.551305 | 13 | 6 |
| | *N. pinetum* | 38.031864 | -84.56518 | 7 | 1 |
| | | 38.032384 | -84.565478 | 15 | 7 |
| | | 38.032407 | -84.568684 | 16 | 25 |
| | | 38.0916 | -84.5602 | 1 | 0 |
| | | 38.0999085 | -84.557832 | 4 | 1 |
| | | 38.210412 | -84.819839 | 1 | 0 |
| | | 38.248397 | -84.54513 | 6 | 0 |
| | | 38.24932 | -84.549175 | 8 | 3 |
| | | 38.250169 | -84.549083 | 1 | 0 |
| | | 38.893228 | -84.557102 | 0 | 2 |
| | | 38.9377 | -84.6335 | 1 | 2 |
| Maryland | *N. lecontei* | 38.499947 | -75.758003 | 2 | 13 |
| Michigan | *N. lecontei* | 43.7596 | -85.7407 | 2 | 0 |
| | | 44.6566667 | -84.695556 | 2 | 0 |
| | | 44.73135 | -84.748931 | 36 | 38 |
| | | 45.5009 | -84.6146 | 0 | 1 |
| | | 45.9262667 | -86.293989 | 39 | 0 |
| | | 45.9485472 | -86.261331 | 6 | 0 |
| | *N. pinetum* | 41.9459 | -85.6507 | 14 | 9 |
| | | 42.257 | -83.7013 | 4 | 11 |
| | | 42.69 | -84.4515 | 2 | 5 |

| | | | | | |
|---|---|---|---|---|---|
| | | 42.9105 | -85.7664 | 4 | 5 |
| | | 44.478233 | -83.836663 | 19 | 6 |
| | | 45.492 | -84.0294 | 6 | 0 |
| North Carolina | *N. lecontei* | 34.693461 | -77.168331 | 13 | 8 |
| | | 35.187 | -81.2913 | 65 | 67 |
| | | 35.7819 | -78.6389 | 1 | 0 |
| | | 35.7835 | -78.8246 | 27 | 30 |
| | | 35.9237 | -79.1032 | 1 | 4 |
| New Hampshire | *N. lecontei* | 42.7553 | -71.2203 | 7 | 2 |
| New York | *N. pinetum* | 43.04105 | -76.45767 | 33 | 15 |
| Ohio | *N. pinetum* | 39.618801 | -83.60369 | 1 | 18 |
| | | 40.7557 | -81.4498 | 3 | 4 |
| Tennessee | *N. pinetum* | 35.8428 | -86.6612 | 0 | 1 |
| | | 36.1156 | -86.6996 | 0 | 2 |
| Virginia | *N. lecontei* | 36.912924 | -75.996487 | 6 | 2 |
| | | 36.916399 | -76.040295 | 3 | 3 |
| | *N. pinetum* | 37.893 | -78.373 | 0 | 1 |
| | | 38.9375511 | -77.281607 | 15 | 5 |
| Wisconsin | *N. lecontei* | 44.1561111 | -90.132222 | 27 | 0 |
| | | 44.9833333 | -88.448056 | 22 | 0 |
| | *N. pinetum* | 43.4288 | -89.4841 | 12 | 13 |

**Table S3. Sampling locations for all *Neodiprion lecontei* and *N. pinetum* females from which eggs were extracted and measured to test for differences in egg size between the species.** *N* represents the number of individual females collected from each sampling site.

| Species | State | Latitude | Longitude | *N* Females | Collection Date |
|---|---|---|---|---|---|
| *N. pinetum* | Kentucky | 38.041545 | -84.441641 | 1 | 6/3/13 |
| *N. pinetum* | Kentucky | 38.032384 | -84.565478 | 1 | 8/19/15 |
| *N. pinetum* | Kentucky | 37.972879 | -84.500398 | 1 | 8/20/15 |
| *N. pinetum* | Kentucky | 37.970609 | -84.497555 | 1 | 8/20/15 |
| *N. pinetum* | Tennessee | 35.98003 | -85.01518 | 1 | 7/11/13 |
| *N. lecontei* | Kentucky | 38.402354 | -85.585931 | 1 | 7/4/15 |
| *N. lecontei* | Kentucky | 38.014 | -84.504 | 2 | 6/9/14 |
| *N. lecontei* | Kentucky | 37.071333 | -84.211417 | 1 | 7/16/13 |
| *N. lecontei* | Tennessee | 35.980274 | -85.015229 | 1 | 6/13/15 |

**Table S4. Sampling locations for each population of *Neodiprion lecontei* and *N. pinetum* used in the mating assays.** The number of colonies (*N*) represents the number of source colonies collected from each sampling site within each population. Typical field caught colonies produce anywhere from 20-150 adults. Adults used in mating assays were either reared directly from these source colonies or were the progeny of these source colonies.

| Cross | Species | Population | Latitude | Longitude | *N* |
|---|---|---|---|---|---|
| 1 | *N. lecontei* | North Carolina | 35.9237 | -79.1032 | 1 |
| | | | 35.7835 | -78.8246 | 6 |
| | | | 35.187 | -81.2913 | 4 |
| | *N. pinetum* | Indiana | 41.3823 | -85.0534 | 1 |
| | | | 41.1655 | -87.2717 | 1 |
| | | | 40.4693 | -86.9132 | 7 |
| | | | 39.1871 | -86.5447 | 1 |
| 2 | *N. lecontei* | Kentucky | 38.1074375 | -84.551305 | 5 |
| | | | 38.09375 | -84.554583 | 2 |
| | | | 38.0146657 | -84.503891 | 10 |
| | *N. pinetum* | Kentucky | 38.9377 | -84.6335 | 2 |
| | | | 38.24932 | -84.549175 | 1 |
| | | | 38.248397 | -84.54513 | 1 |
| | | | 38.0324069 | -84.568684 | 5 |
| 3 | *N. lecontei* | Kentucky | 38.1074375 | -84.551305 | 3 |
| | | | 38.09375 | -84.554583 | 3 |
| | | | 38.0146657 | -84.503891 | 12 |
| | | | 37.9868001 | -84.416332 | 1 |
| | *N. pinetum* | Michigan | 45.492 | -84.0294 | 1 |
| | | | 42.9105 | -85.7664 | 1 |
| | | | 42.69 | -84.4515 | 2 |
| | | | 42.257 | -83.7013 | 1 |
| | | | 41.9459 | -85.6507 | 2 |

**Table S5. Type III ANOVA table for linear regression model for pine needle width.** Significant *P*-values ($P < 0.05$) are indicated in bold.

| Factor | Sum Sq | df | F value | *P*-value |
|---|---|---|---|---|
| (Intercept) | 0.019 | 1 | 1.44 | 0.23 |
| latitude | 0.012 | 1 | 0.91 | 0.34 |
| host species | 0.50 | 10 | 3.76 | **0.000057** |
| latitude*host species | 0.57 | 10 | 4.26 | **0.0000084** |
| Residuals | 11.43 | 856 | | |

**Table S6. Post-hoc estimated marginal means for comparisons of needle width between all pairs of 11 *Pinus* pine species: *P. taeda* (loblolly), *P. palustris* (longleaf), *P. echinata* (shortleaf), *P. elliottii* (slash), *P. clausa* (sand), *P. glabra* (spruce), *P. virginiana* (virginia), *P. rigida* (pitch), *P. resinosa* (red), *P. banksiana* (jack), and *P. strobus* (white).** *P*-values are corrected for multiple testing using the false discovery rate (FDR). Significant *P*-values ($P < 0.05$) are indicated in bold.

| Comparison | Estimate | SE | df | t-ratio | *P*-value |
|---|---|---|---|---|---|
| jack - loblolly | 0.08125 | 0.1341 | 856 | 0.606 | 0.6114 |
| jack - longleaf | -0.00779 | 0.1449 | 856 | -0.054 | 0.9571 |
| jack - pitch | 0.14002 | 0.1358 | 856 | 1.031 | 0.3776 |
| jack - red | -0.25163 | 0.2472 | 856 | -1.018 | 0.3776 |
| jack - sand | 0.77088 | 0.5075 | 856 | 1.519 | 0.2248 |
| jack - shortleaf | 0.20591 | 0.1361 | 856 | 1.512 | 0.2248 |
| jack - slash | -0.07757 | 0.175 | 856 | -0.443 | 0.6955 |
| jack - spruce | 0.27007 | 0.174 | 856 | 1.552 | 0.2217 |
| jack - virginia | 0.04784 | 0.1344 | 856 | 0.356 | 0.7491 |
| jack - white | 0.65046 | 0.1354 | 856 | 4.803 | **< 0.0001** |
| loblolly - longleaf | -0.08904 | 0.0604 | 856 | -1.474 | 0.228 |
| loblolly - pitch | 0.05878 | 0.0333 | 856 | 1.766 | 0.1782 |
| loblolly - red | -0.33287 | 0.2092 | 856 | -1.591 | 0.2176 |
| loblolly - sand | 0.68964 | 0.4901 | 856 | 1.407 | 0.2441 |
| loblolly - shortleaf | 0.12466 | 0.0347 | 856 | 3.593 | **0.0019** |
| loblolly - slash | -0.15882 | 0.1152 | 856 | -1.378 | 0.2505 |
| loblolly - spruce | 0.18882 | 0.1137 | 856 | 1.66 | 0.2056 |
| loblolly - virginia | -0.0334 | 0.0269 | 856 | -1.241 | 0.3034 |
| loblolly - white | 0.56921 | 0.0318 | 856 | 17.89 | **< 0.0001** |
| longleaf - pitch | 0.14782 | 0.0642 | 856 | 2.304 | 0.0787 |
| longleaf - red | -0.24383 | 0.2163 | 856 | -1.127 | 0.3425 |
| longleaf - sand | 0.77868 | 0.4932 | 856 | 1.579 | 0.2176 |
| longleaf - shortleaf | 0.2137 | 0.0649 | 856 | 3.293 | **0.0047** |
| longleaf - slash | -0.06978 | 0.1276 | 856 | -0.547 | 0.6305 |
| longleaf - spruce | 0.27786 | 0.1263 | 856 | 2.201 | 0.087 |
| longleaf - virginia | 0.05564 | 0.0611 | 856 | 0.91 | 0.4246 |
| longleaf - white | 0.65825 | 0.0634 | 856 | 10.38 | **< 0.0001** |
| pitch - red | -0.39165 | 0.2103 | 856 | -1.862 | 0.1525 |
| pitch - sand | 0.63086 | 0.4906 | 856 | 1.286 | 0.2877 |
| pitch - shortleaf | 0.06589 | 0.0409 | 856 | 1.612 | 0.2176 |
| pitch - slash | -0.2176 | 0.1172 | 856 | -1.856 | 0.1525 |
| pitch - spruce | 0.13005 | 0.1158 | 856 | 1.123 | 0.3425 |
| pitch - virginia | -0.09218 | 0.0345 | 856 | -2.671 | **0.0326** |

| | | | | | |
|---|---|---|---|---|---|
| pitch - white | 0.51044 | 0.0384 | 856 | 13.276 | **< 0.0001** |
| red - sand | 1.02251 | 0.5323 | 856 | 1.921 | 0.1442 |
| red - shortleaf | 0.45754 | 0.2105 | 856 | 2.173 | 0.087 |
| red - slash | 0.17405 | 0.2375 | 856 | 0.733 | 0.5315 |
| red - spruce | 0.52169 | 0.2368 | 856 | 2.203 | 0.087 |
| red - virginia | 0.29947 | 0.2094 | 856 | 1.43 | 0.2405 |
| red - white | 0.90208 | 0.2101 | 856 | 4.294 | **0.0001** |
| sand - shortleaf | -0.56497 | 0.4907 | 856 | -1.151 | 0.3425 |
| sand - slash | -0.84845 | 0.5028 | 856 | -1.687 | 0.2022 |
| sand - spruce | -0.50081 | 0.5025 | 856 | -0.997 | 0.3817 |
| sand - virginia | -0.72304 | 0.4902 | 856 | -1.475 | 0.228 |
| sand - white | -0.12042 | 0.4905 | 856 | -0.246 | 0.821 |
| shortleaf - slash | -0.28348 | 0.1176 | 856 | -2.41 | 0.0636 |
| shortleaf - spruce | 0.06416 | 0.1162 | 856 | 0.552 | 0.6305 |
| shortleaf - virginia | -0.15807 | 0.0359 | 856 | -4.405 | **0.0001** |
| shortleaf - white | 0.44455 | 0.0397 | 856 | 11.203 | **< 0.0001** |
| slash - spruce | 0.34764 | 0.1599 | 856 | 2.174 | 0.087 |
| slash - virginia | 0.12542 | 0.1156 | 856 | 1.085 | 0.3559 |
| slash - white | 0.72803 | 0.1168 | 856 | 6.232 | **< 0.0001** |
| spruce - virginia | -0.22223 | 0.1141 | 856 | -1.948 | 0.1424 |
| spruce - white | 0.38039 | 0.1153 | 856 | 3.298 | **0.0047** |
| virginia - white | 0.60262 | 0.0331 | 856 | 18.205 | **< 0.0001** |

**Table S7. Type III ANOVA table for linear regression model for adult sawfly body size.** Significant *P*-values ($P < 0.05$) are indicated in bold.

| **Factor** | **Sum Sq** | **df** | **F value** | ***P*-value** |
|---|---|---|---|---|
| (Intercept) | 299.57 | 1 | 883.96 | **$< 2.2 \times 10^{-16}$** |
| latitude | 12.06 | 1 | 35.57 | **$3.34 \times 10^{-09}$** |
| species | 9.97 | 1 | 29.43 | **$7.18 \times 10^{-08}$** |
| sex | 19.32 | 1 | 57.0076 | **$9.26 \times 10^{-14}$** |
| latitude*species | 14.31 | 1 | 42.22 | **$1.25 \times 10^{-10}$** |
| latitude*sex | 1.72 | 1 | 5.08 | **0.024** |
| species*sex | 3.53 | 1 | 10.41 | **0.0013** |
| Residuals | 363.3 | 1072 | | |

**Table S8. Type III ANOVA table for logistic regression model for hatching success (proportion of eggs that hatched) modeled as a function of egg area and host (*Pinus banksiana* or *P. strobus*).** Significant *P*-values ($P < 0.05$) are indicated in bold.

| **Factor** | **Chi-square** | **df** | ***P*-value** |
|---|---:|:---:|---:|
| egg area | 19.60 | 1 | **0.0000095** |
| host | 8.31 | 1 | **0.0039** |
| egg area*host | 11.61 | 1 | **0.00066** |

**Table S9. The strength of prezygotic isolation between *Neodiprion lecontei* and *N. pinetum* varies among crosses from different geographic regions (A) and among different pair types within each cross and globally (all crosses combined) (B).** Reproductive isolation strength, after Sobel and Chen (2014), is abbreviated as RI. *P*-values represent the results of the one-tailed Fisher's exact tests (A) and the results of pairwise tests for differences in mating outcomes between the four pair types, after FDR correction for multiple testing. Significant *P*-values ($P < 0.05$) are indicated in bold. Cross 1 is North Carolina *N. lecontei* x Indiana *N. pinetum*; Cross 2 is Kentucky *N. lecontei* x Kentucky *N. pinetum*; Cross 3 is Kentucky *N. lecontei* x Michigan *N. pinetum*.

(A) Strength of prezygotic isolation for each cross and globally (all crosses combined)

| **Cross Number** | ***N. lecontei* Source** | ***N. pinetum* Source** | **RI** | ***P*-value** | **Odds Ratio** |
|---|---|---|---|---|---|
| 1 | North Carolina | Indiana | 0.19 | 0.26 | 1.46 |
| 2 | Kentucky | Kentucky | 0.37 | **0.047** | 3.00 |
| 3 | Kentucky | Michigan | 0.77 | **0.00019** | 13.37 |
| Global | all above | all above | 0.39 | **0.00038** | 2.86 |

(B) FDR-corrected *P*-values for mating outcomes between each of the four pair types

| **Cross Number** | **Pair types** | *N. lecontei* ♀ *N. lecontei* ♂ | *N. lecontei* ♀ *N. pinetum* ♂ | *N. pinetum* ♀ *N. lecontei* ♂ |
|---|---|---|---|---|
| 1 | | | | |
| | *N. lecontei* ♀ *N. pinetum* ♂ | **0.00098** | | |
| | *N. pinetum* ♀ *N. lecontei* ♂ | 1 | **0.0012** | |
| | *N. pinetum* ♀ *N. pinetum* ♂ | **0.036** | 0.29 | 0.052 |
| 2 | | | | |
| | *N. lecontei* ♀ *N. pinetum* ♂ | 0.64 | | |
| | *N. pinetum* ♀ *N. lecontei* ♂ | 0.64 | 1 | |
| | *N. pinetum* ♀ *N. pinetum* ♂ | 1 | 0.64 | 0.64 |

| 3 | *N. lecontei* ♀ *N. lecontei* ♂ | *N. lecontei* ♀ *N. pinetum* ♂ | *N. pinetum* ♀ *N. lecontei* ♂ |
| --- | --- | --- | --- |
| *N. lecontei* ♀ *N. pinetum* ♂ | **0.0051** | | |
| *N. pinetum* ♀ *N. lecontei* ♂ | **0.0051** | 1 | |
| *N. pinetum* ♀ *N. pinetum* ♂ | 0.20 | 0.20 | 0.20 |

| Global | *N. lecontei* ♀ *N. lecontei* ♂ | *N. lecontei* ♀ *N. pinetum* ♂ | *N. pinetum* ♀ *N. lecontei* ♂ |
| --- | --- | --- | --- |
| *N. lecontei* ♀ *N. pinetum* ♂ | **0.0000071** | | |
| *N. pinetum* ♀ *N. lecontei* ♂ | **0.030** | **0.023** | |
| *N. pinetum* ♀ *N. pinetum* ♂ | **0.030** | **0.023** | 1 |

**Table S10. Type III ANOVA tables for logistic regression model for binary mating outcome modeled as a function of female species, male species, female species x male species interaction, and size differential (female body length – male body length).** Cross 1 is North Carolina *Neodiprion lecontei* x Indiana *N. pinetum*; Cross 2 is Kentucky *N. lecontei* x Kentucky *N. pinetum*; Cross 3 is Kentucky *N. lecontei* x Michigan *N. pinetum*. For the global model, "cross" was included as a random effect. Significant *P*-values ($P < 0.05$) are indicated in bold.

| Cross | Factor | Chi-square | df | *P*-value |
|---|---|---|---|---|
| 1 | Female species | 0.48 | 1 | 0.49 |
| | Male species | 14.40 | 1 | **0.00015** |
| | Size differential | 15.13 | 1 | **0.0001** |
| | Female species x male species | 1.86 | 1 | 0.17 |
| 2 | Female species | 0.13 | 1 | 0.71 |
| | Male species | 0.06 | 1 | 0.81 |
| | Size differential | 14.78 | 1 | **0.00012** |
| | Female species x male species | 0.82 | 1 | 0.36 |
| 3 | Female species | 2.70 | 1 | 0.10 |
| | Male species | 1.39 | 1 | 0.24 |
| | Size differential | 9.57 | 1 | **0.0020** |
| | Female species x male species | 3.19 | 1 | 0.074 |
| Global | Female species | 1.43 | 1 | 0.23 |
| | Male species | 8.34 | 1 | **0.0039** |
| | Size differential | 30.79 | 1 | **$2.87 \times 10^{-08}$** |
| | Female species x male species | 3.81 | 1 | 0.051 |

**Table S11. Type III ANOVA tables for logistic regression model for binary male mating attempts modeled as a function of female species, male species, female species x male species interaction, and size differential (female body length – male body length).** Cross 1 is North Carolina *Neodiprion lecontei* x Indiana *N. pinetum*; Cross 2 is Kentucky *N. lecontei* x Kentucky *N. pinetum*; Cross 3 is Kentucky *N. lecontei* x Michigan *N. pinetum*. For the global model, "cross" was included as a random effect. Significant *P*-values ($P < 0.05$) are indicated in bold.

| Cross | Factor | Chi-square | df | *P*-value |
|---|---|---|---|---|
| 1 | Female species | 0.059 | 1 | 0.81 |
| | Male species | 10.86 | 1 | **0.00098** |
| | Size differential | 3.71 | 1 | 0.054 |
| | Female species x male species | 0.27 | 1 | 0.60 |
| 2 | Female species | 1.63 | 1 | 0.20 |
| | Male species | 0.59 | 1 | 0.44 |
| | Size differential | 0.16 | 1 | 0.69 |
| | Female species x male species | 0.19 | 1 | 0.66 |
| 3 | Female species | 0.00001 | 1 | 0.998 |
| | Male species | 0.55 | 1 | 0.46 |
| | Size differential | 2.68 | 1 | 0.10 |
| | Female species x male species | 0.81 | 1 | 0.37 |
| Global | Female species | 0.035 | 1 | 0.85 |
| | Male species | 10.53 | 1 | **0.0012** |
| | Size differential | 5.26 | 1 | **0.022** |
| | Female species x male species | 1.04 | 1 | 0.31 |

**Table S12. Results of one-sided t-tests testing whether heterospecific pairs had significantly greater size differentials than conspecific pairs.** Results are shown for each cross and for the global analysis. Cross 1 is North Carolina *Neodiprion lecontei* x Indiana *N. pinetum*; Cross 2 is Kentucky *N. lecontei* x Kentucky *N. pinetum*; Cross 3 is Kentucky *N. lecontei* x Michigan *N. pinetum*. Significant *P*-values ($P < 0.05$) are indicated in bold.

| Cross | t value | df | *P*-value |
|---|---|---|---|
| 1 | 0.76 | 138 | 0.22 |
| 2 | 1.34 | 55 | 0.093 |
| 3 | 2.56 | 42 | **0.0070** |
| Global | 2.41 | 243 | **0.0084** |

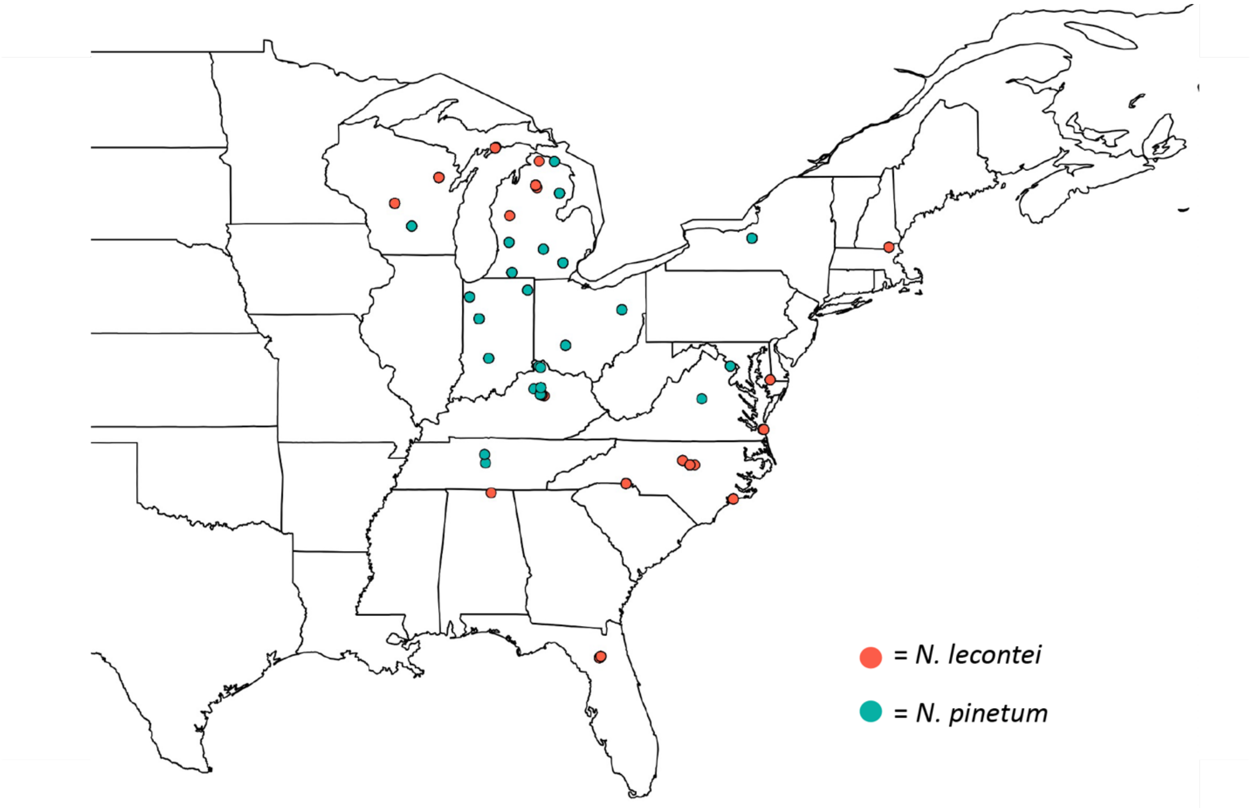


**Figure S1. Map depicting the sampling locations of all *Neodiprion lecontei* (coral circles) and *N. pinetum* (teal circles) individuals measured to characterize geographic variation in body length.**

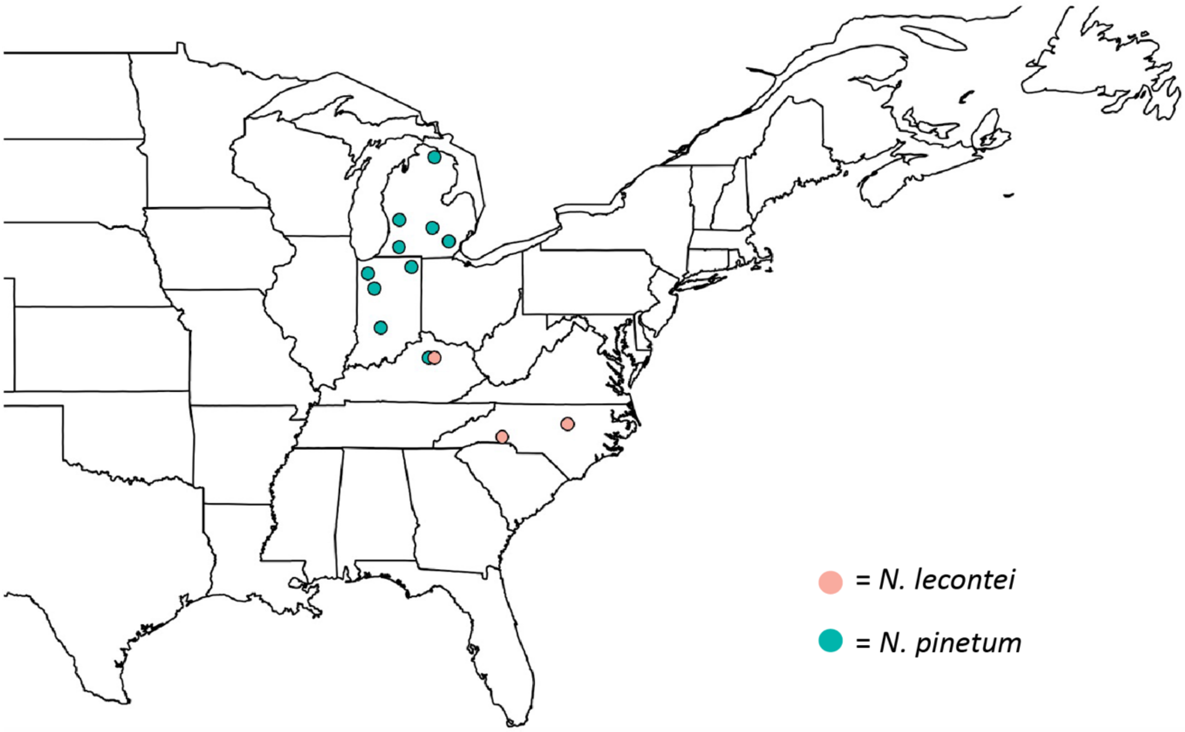


**Figure S2. Map depicting the sampling locations of all *Neodiprion lecontei* colonies (coral circles) and *N. pinetum* colonies (teal circles) used in the mating assays conducted for this study.** Adult females and males from these colonies were either used directly in mating assays or used to propagate a first lab generation (colonies were grouped by state), from which adults were then used in mating assays.

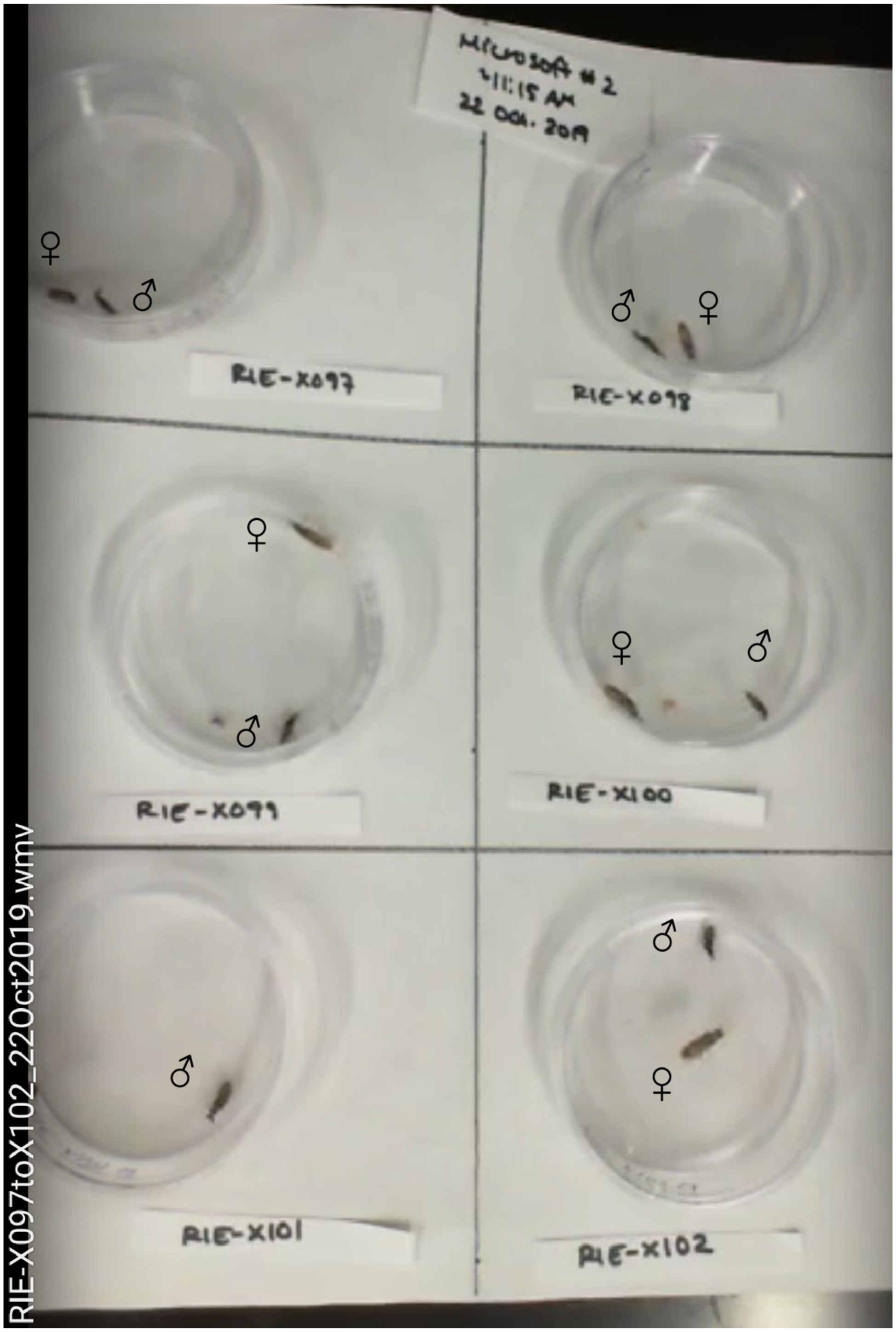


**Figure S3. Set-up of each arena in no-choice mating assays.** Each arena was divided into 6 equally sized sections, and a small petri dish containing a single male and female were placed in each section. Each pair was assigned an arbitrary identifier to avoid biases during recording of mating outcome and male mating attempts.

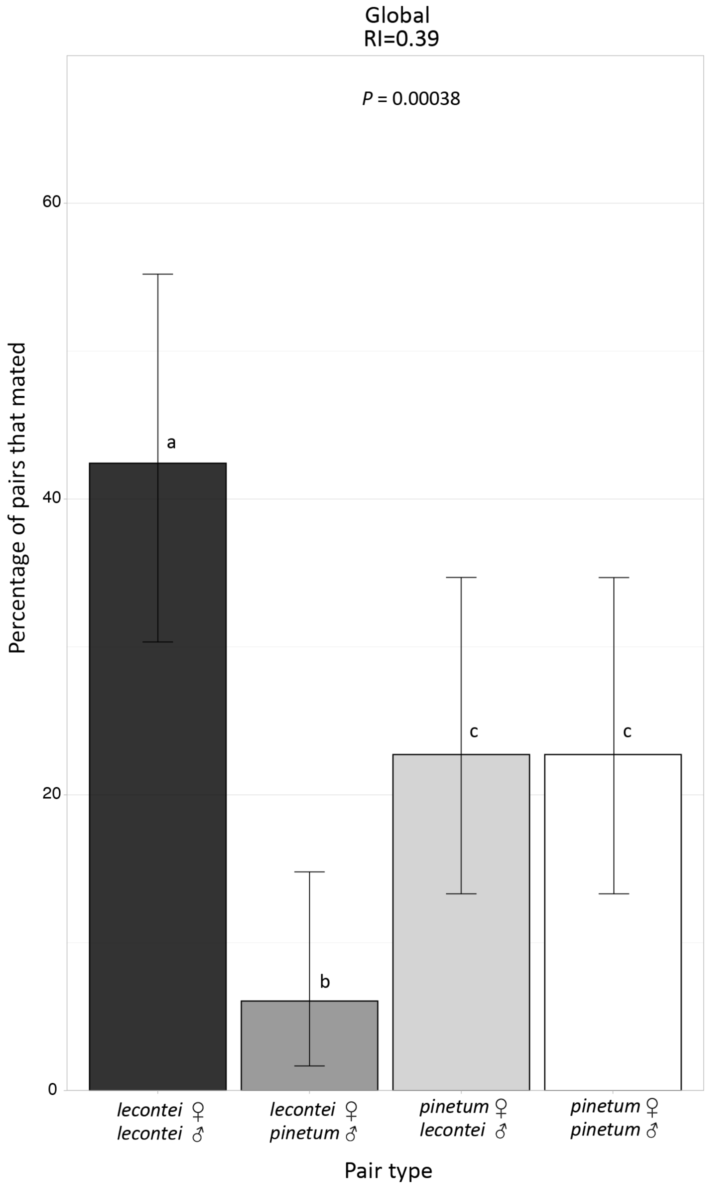


**Figure S4. Prezygotic isolation between *Neodiprion lecontei* and *N. pinetum*.** The percentage of pairs that mated as a function of pair type for all crosses combined (global) are presented. The reproductive isolation (RI) value represents the strength of reproductive isolation between *N. lecontei* and *N. pinetum* as calculated after Sobel and Chen (2014). The *P*-value listed indicates the result of the one-sided Fisher's exact test; a significant *P*-value indicates that conspecific pairs mated more often than heterospecific pairs. Error bars represent the 95% Clopper-Pearson confidence intervals for the percentage of pairs that mated. Letters indicate pair types that differ significantly at $P < 0.05$, after FDR correction for multiple testing (Table S9). Shared letters indicate pair types that were not significantly different.

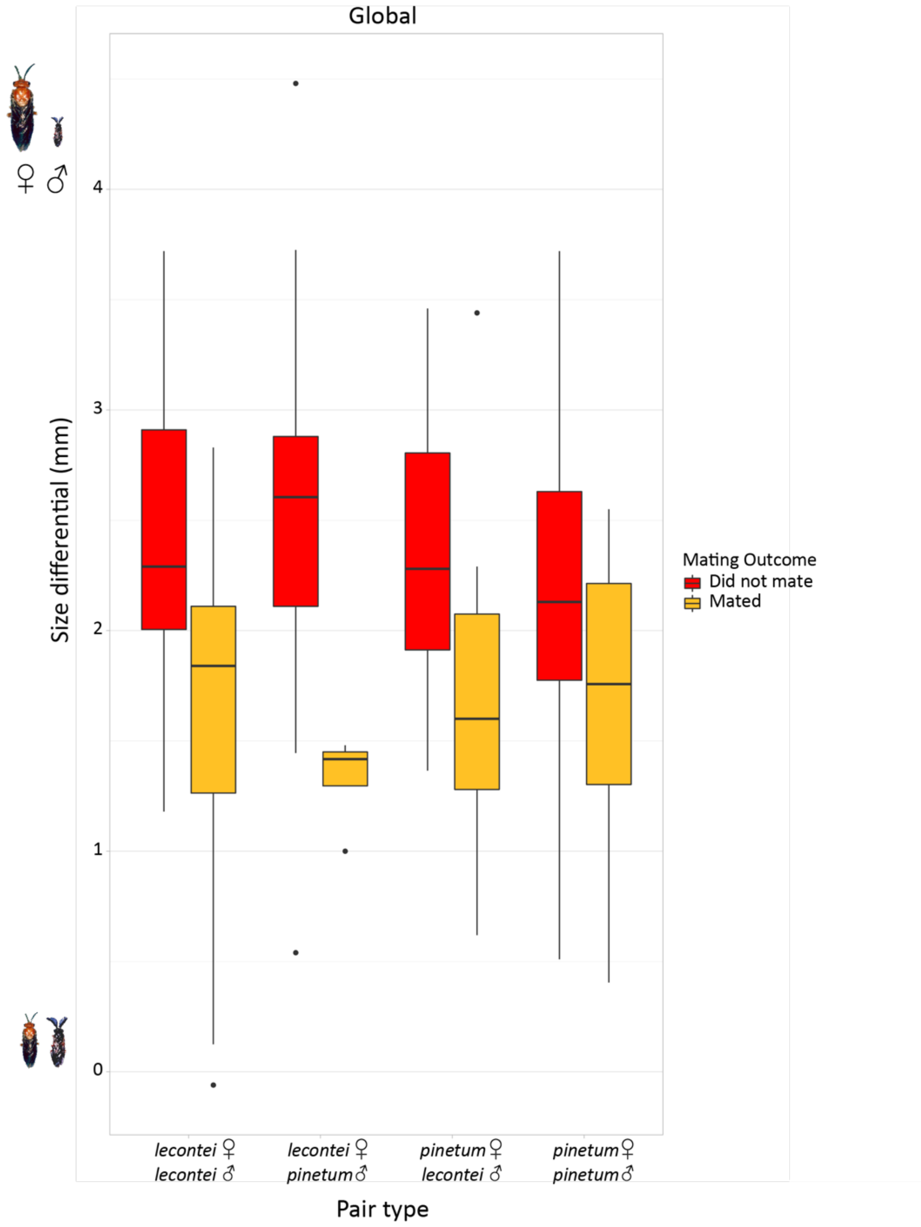


**Figure S5. Across all pair types when all crosses are combined (Global), pairs that did not mate (red boxes) had greater size differentials than pairs that did mate (yellow boxes).** Size differentials are female body length minus male body length; higher values reflect larger size differences (adults not drawn to scale). Boxes represent interquartile ranges (median ± 2 SD), with size differential outliers indicated as points. Note that while we are showing size differentials on the y-axis to visualize the consistency of this pattern, mating outcome was modeled as the response variable. Size differential had a significant effect on mating outcome globally (Table S10).

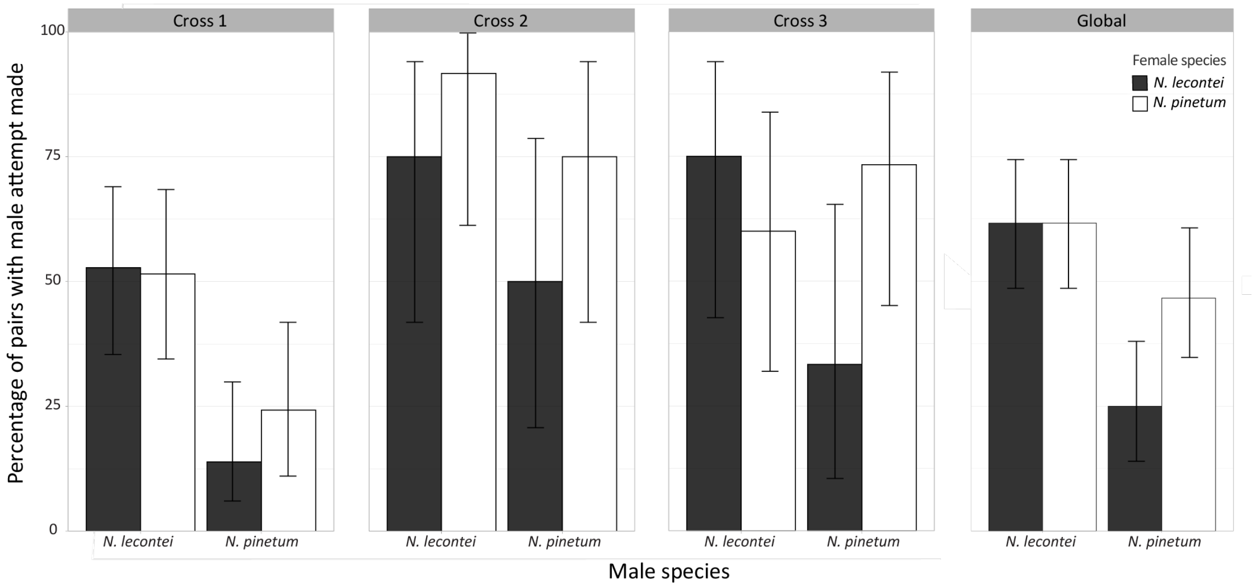


**Figure S6. Variation in male willingness/motivation to mate.** Each panel gives the percentage of pairs that a male attempt to mate was made as a function of pair type for one of the three crosses and globally (all crosses combined): Cross 1 is North Carolina *Neodiprion lecontei* x Indiana *N. pinetum*; Cross 2 is Kentucky *N. lecontei* x Kentucky *N. pinetum*; Cross 3 is Kentucky *N. lecontei* x Michigan *N. pinetum*. Error bars represent the 95% Clopper-Pearson confidence intervals for the percentage of pairs that a male attempt to mate was made.

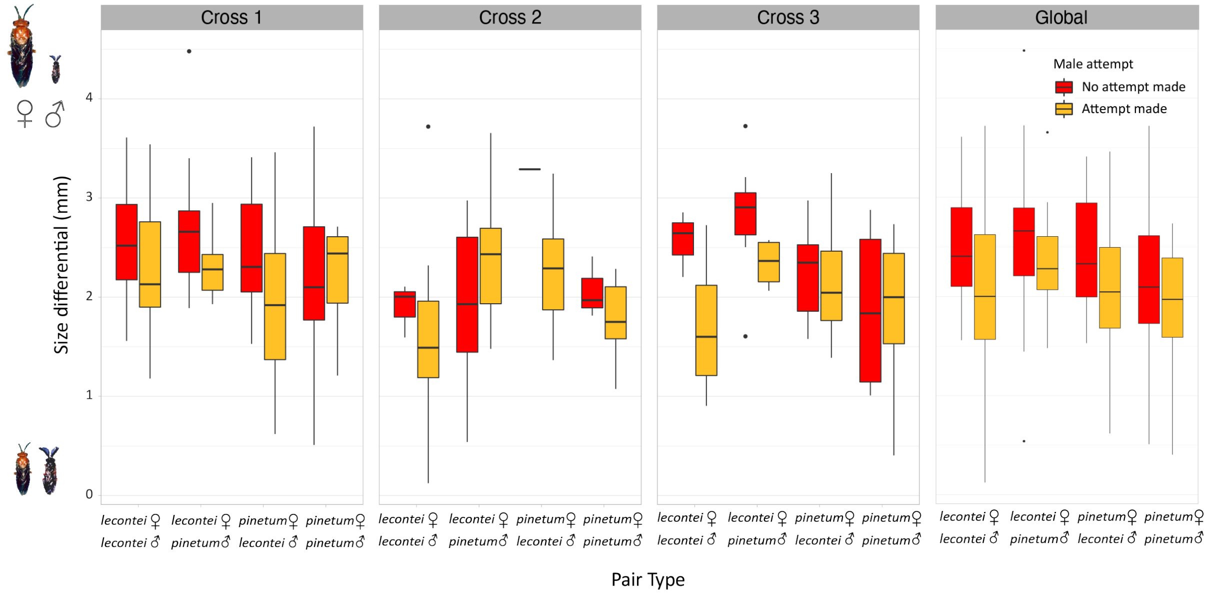


**Figure S7. Across all pair types, pairs where the males did not make an attempt to mate (red boxes) had greater size differentials than pairs where the male made at least one attempt to mate (yellow boxes).** Cross 1 is North Carolina *Neodiprion lecontei* x Indiana *N. pinetum*; Cross 2 is Kentucky *N. lecontei* x Kentucky *N. pinetum*; Cross 3 is Kentucky *N. lecontei* x Michigan *N. pinetum*; global is all crosses combined. Size differentials are female body length minus male body length; higher values reflect larger size differences (adults not drawn to scale). Boxes represent interquartile ranges (median ± 2 SD), with size differential outliers indicated as points. Note that while we are showing size differentials on the y-axis to visualize the consistency of this pattern, male attempts was modeled as the binary response variable. Size differential had a significant effect on male attempts in the global analysis only (Table S11).

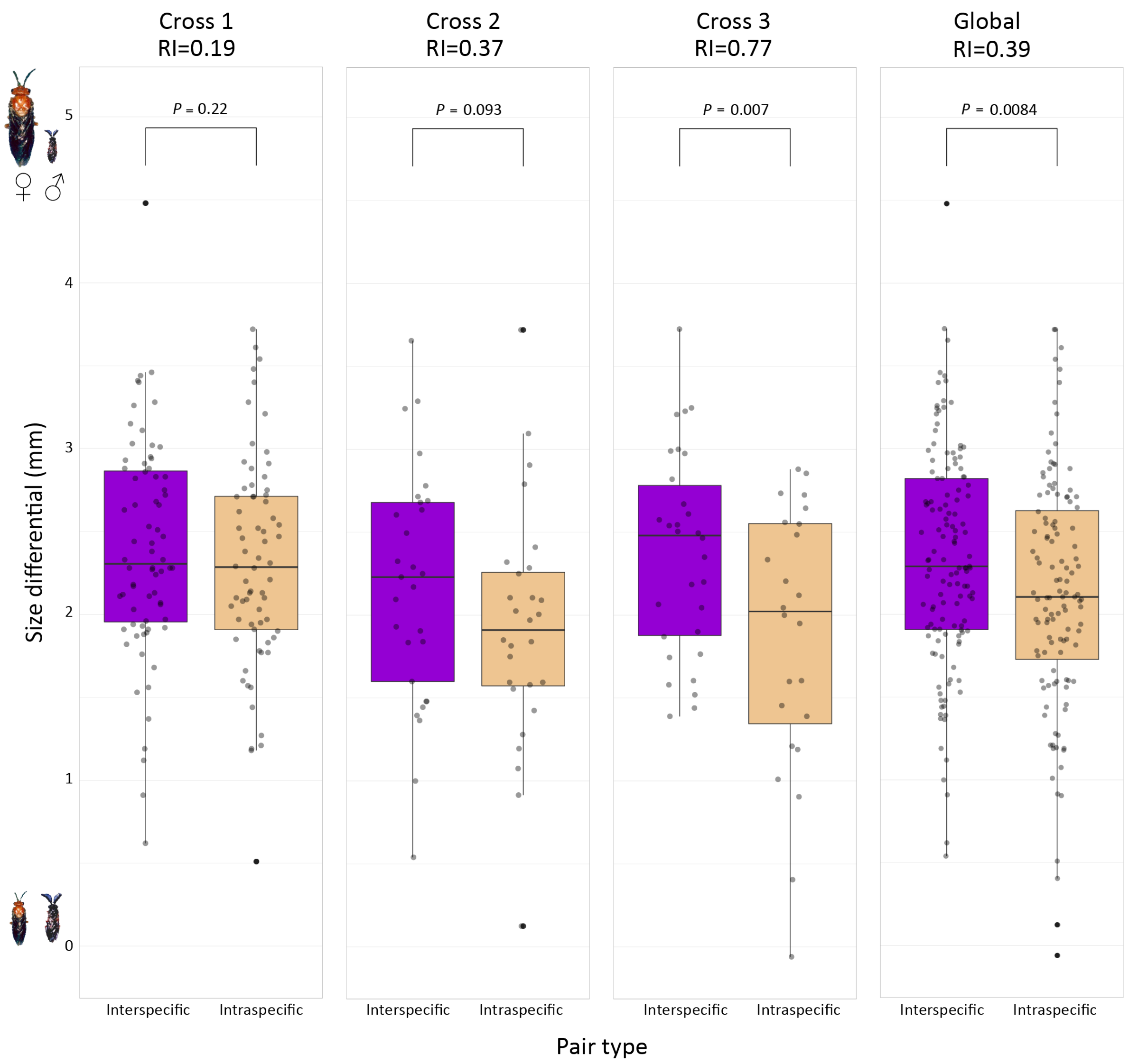


**Figure S8. Size differentials (in mm) for interspecific pairs (purple boxes) compared to intraspecific pairs (tan boxes) for each cross and globally (all crosses combined).** Cross 1 is North Carolina *Neodiprion lecontei* x Indiana *N. pinetum*; Cross 2 is Kentucky *N. lecontei* x Kentucky *N. pinetum*; Cross 3 is Kentucky *N. lecontei* x Michigan *N. pinetum*. Size differentials are female body length minus male body length; higher values reflect larger size differences (adults not drawn to scale). Boxes represent interquartile ranges (median ± 2 SD), with size differential outliers indicated as black points. Gray points represent the raw data points. The strength of prezygotic isolation (RI) for each cross and globally (all crosses combined) were calculated after Sobel and Chen (2014). *P*-values (*P*) were calculated via one-sided t-tests to evaluate whether the size differential for interspecific pairs was significantly greater than intraspecific pairs (Table S12).

**Video S1. A successful mating between a *Neodiprion lecontei* female and male.**
